# Tau seeding and spreading in vivo is supported by both AD-derived fibrillar and oligomeric tau

**DOI:** 10.1101/2023.03.28.534418

**Authors:** Anastasie Mate de Gerando, Lindsay A Welikovitch, Anita Khasnavis, Caitlin Commins, Calina Glynn, Joshua E Chun, Romain Perbet, Bradley T Hyman

## Abstract

Insoluble fibrillar tau, the primary constituent of neurofibrillary tangles, has traditionally been thought to be the biologically active, toxic form of tau mediating neurodegeneration in Alzheimer’s disease. More recent studies have implicated soluble oligomeric tau species, referred to as high molecular weight (HMW) due to its properties on size exclusion chromatography, in tau propagation across neural systems. These two forms of tau have never been directly compared. We prepared sarkosyl insoluble and HMW tau from the frontal cortex of Alzheimer patients and compared their properties using a variety of biophysical and bioactivity assays. Sarkosyl insoluble fibrillar tau is comprised of abundant paired helical filaments (PHF) as quantified by electron microscopy (EM), and is more resistant to proteinase K, compared to HMW tau which is mostly in an oligomeric form. Sarkosyl insoluble and HMW tau are nearly equivalent in potency in a HEK cell bioactivity assay for seeding aggregates and their injection reveals similar local uptake into hippocampal neurons in PS19 Tau transgenic mice. However, the HMW preparation appears to be far more potent in inducing a glial response including Clec7a-positive rod-microglia in the absence of neurodegeneration or synapse loss and promotes more rapid propagation of misfolded tau to distal, anatomically connected regions, such as entorhinal and perirhinal cortices. These data suggest that soluble HMW tau has similar properties to fibrillar sarkosyl insoluble tau with regard to tau seeding potential but may be equal or even more bioactive with respect to propagation across neural systems and activation of glial responses, both relevant tau-related Alzheimer phenotypes.

## Introduction

Alzheimer’s disease (AD) is a devastating neurological disorder resulting in progressive cognitive decline and memory loss. Since their initial observation [1], fibrillar tau protein inclusions in neurons, known as neurofibrillary tangles (NFT), have been associated with disease progression and severity [2, 19]. However, whether NFTs are the entire story has been questioned given that transgenic mouse models of tauopathy develop neurodegeneration, gliosis, and synaptic loss before the appearance of NFTs [45, 53, 66]. The hypothesis of the existence of a toxic, non-fibrillar oligomeric tau species has thus emerged [7, 13, 43, 62]. In addition, the rate of clinical progression in AD patients is tightly associated with the properties of soluble oligomeric HMW tau [17], as well as analogous differences in fibrillar tau [32], raising the question of how fibrillar and oligomeric tau compare.

Tau fibrils [3, 8, 20, 22] or tau oligomers [17, 36, 57] extracted from AD human brain support native tau recruitment and aggregation, termed as seeding, in cellular and mouse models. Although different laboratories have studied both tau preparations, a direct comparison of tau fibrils and oligomers derived from individual AD brains has yet to be performed. We directly compared the biochemical and cellular properties of fibrillar and oligomeric tau to decipher their respective contributions to AD pathogenesis. From each AD case (n=8), we isolated Tau fibrils (SARK tau) and oligomers (HMW tau) by sarkosyl extraction [42] and size-exclusion chromatography [57] respectively. While both tau species can seed and induce similar levels of tau pathology *in vitro* and *in vivo*, they trigger differential glial responses. In addition, it appears that HMW oligomeric species are more potent at propagating across neural systems to seed distant, anatomically connected sites. These results highlight the potential importance of non-fibrillar oligomeric tau species in AD pathogenesis and suggest that some therapeutic approaches targeting fibrillar conformations may miss the equally (or perhaps more) bioactive HMW oligomeric species.

## Materials and methods

### Human tissue

Human participants with AD were selected from the Massachusetts Alzheimer’s Disease Research Center brain bank using the following criteria: (i) clinical diagnosis of dementia due to probable AD; (ii) postmortem confirmation of AD diagnosis; (iii) Braak stage V-VI determined by total tau immunostain and Bielchowsky’s silver stain of NFTs; (iv) minimal comorbidities; and (v) considered as “high seeders” according to our previously published paper [17]. Human brain tissues were collected with informed consent of patients or their relatives and approval of local institutional review boards at Massachusetts General Hospital. Approximatively 5 g of frozen frontal cortex corresponding to Broadman Area 8/9 and 7 (BA 8/9 and BA 7) from 8 different AD cases (age-matched male and female donors) were dissected and kept at -80°C until processing for either SARK or HMW tau extraction.

### Sarkosyl tau extraction

For each case, frozen human tissue (5 g/case) was thawed on wet ice, the meninges and white matter were removed and the grey matter processed according to previously described sarkosyl extraction protocols [22, 42]. Grey matter was Dounce homogenized in 9 v/w high salt buffer (10mM Tris pH 7.4, 10% sucrose, 0.8 M NaCl, 1 mM EDTA, 0.1% sarkosyl and 1X protease/phosphatase inhibitor cocktail, #5872, Cell signaling) in a 15 ml glass homogenizer with 15 up and down strokes at 70% power by hand on ice. Homogenates were transferred to a 50 ml Falcon tube and centrifuged 10,000 g for 10 min at 4°C. The supernatants were collected and filtered through a kimwipe into a 50 ml Falcon tube. Homogenization and centrifugation/filtration steps were repeated once to increase the yield. A solution of 25% sarkosyl in distilled water was added to the total volumes of supernatant to get a final concentration of 1% sarkosyl and left incubating for 1 hour under agitation at room temperature (RT). Supernatants were then ultracentrifuged at 300,000 g for 1 hour at 4°C, the myelin and supernatant carefully discarded, and the pellets rinsed twice and resuspended in 1 ml PBS before ultracentrifugation at 250,000 g for 30 min at 4°C. Pellets were resuspended and broken down into small pieces in 1 ml PBS, and left overnight under agitation at RT. After a quick spin (1 min at 1,000 g), pellets were broken up using a 27-gauge needle and sonicated with 20 short pulses at power 2 on ice using a hand-held sonicator (QSonica). Samples were then centrifuged 30 min at 100,000 g at 4°C, the supernatants discarded, the pellets resuspended in 200 µl PBS 1X and sonicated 60 short pulses before centrifugation 30 min at 10,000 g at 4°C. The final supernatant contained the sarkosyl-insoluble tau species (SARK tau) and were stored at -80°C until further use. Depending on the experimental condition, SARK tau samples were extemporaneously sonicated for 60 short pulses on power 2 on ice using a hand-held sonicator as recommended in the literature [22, 42].

### HMW tau extraction

For each case, frozen human tissue (5 g/case) was dissected as described above, and the grey matter was Dounce homogenized in 5 volumes per weight (v/w) buffer containing phosphate buffer saline (PBS) and 1X protease inhibitor (#5871, Cell signaling) in a 15 ml glass homogenizer with 30 up and down strokes at 70% power by hand on ice. Homogenates were transferred to a 50 ml Falcon tube and centrifuged at 10,000 g for 10 min at 4°C. The supernatants were collected and kept at - 80°C until further processing. The total volumes of soluble brain extracts were separated by SEC as previously described [17] on a single Superdex200 10/300GL column (no. 17-5175-01, GE Healthcare) in PBS (no. P3813, Sigma-Aldrich, filtered through a 0.2-um membrane filter), at a flow rate of 0.5 ml/min using an AKTA purifier 10 (GE Healthcare). For each run, 5 ml of soluble brain extract was loaded onto the column and 28 fractions of 2.5 ml were collected. Fractions 2, 3 and 4 containing HMW tau (400-600 kDa) were pooled and centrifuged at 150,000 g for 30 min at 4°C. Concentrated pellets were subsequently resuspended in PBS 1X and stored at -80°C until further use.

### Tau immunodepletion

To verify the tau-specific effects of the SARK and HMW samples, we immunodepleted total tau using HT7 antibody (MN1000, Invitrogen). PureProteome Protein G Magnetic Beads (LSKMAGG02, Millipore Corp) were resuspended by vortexing and 25 µl of the suspended beads were used for each sample. Using a magnetic rack, the storage buffer was removed and discarded. Beads were washed three times with 50 µl PBS 1X, and then 50 µl of HT7 antibody was added. The bead-HT7 solution was incubated for 1 hour at 4°C with head-over-tail rotation. The antibody flow-through supernatant was removed using a magnetic rack, and the beads were washed two more times with PBS. Next, 75 µl of either SARK or HMW tau were incubated with antibody-bead complexes overnight at 4°C with head-over-tail rotation. The next morning, tau-immunodepleted supernatant was collected using a magnetic rack and stored at -80°C until further use. Immunodepletion was repeated twice more on the collected supernatant to fully capture residual tau in the sample. Tau immunodepletion was confirmed by western blotting and tau seeding assay normalizing on total protein levels (Supplementary Fig. 4a-c).

### Total tau quantification by Western blot

Each SARK and HMW tau sample was run on a denaturing Western blot (WB) to quantify total tau monomer equivalents. Samples were diluted in 1X NuPAGE LDS sample buffer (ThermoFisher) and 1X NuPAGE sample reducing agent (ThermoFisher), incubated for 5 min at 95°C and loaded onto a NuPAGE 4-12% Tris/Bis gel (ThermoFisher) with NuPAGE MOPS running buffer (ThermoFisher). Proteins were transferred onto a nitrocellulose membrane using iBlot 2 gel transfer device (7 min at 25 V, ThermoFisher). Membranes were incubated for 1 hour at room temperature (RT) with agitation in Intercept blocking buffer (Licor), then incubated overnight at 4°C with anti-total tau primary antibody solution (1: 5000, DAKO A0024). After 1 hour incubation at RT in the corresponding secondary antibody solution (donkey anti-rabbit 800), membranes were revealed using Licor Odyssey Clx. Recombinant 2N4R human Tau (Tau441, AG960, Millipore) at serial dilutions ranging from 20 µg/ml to 0 µg/ml total tau was used to generate a calibration curve and calculate total tau in our samples.

### Total tau quantification by ELISA

Each SARK and HMW tau sample was also run on an MSD ELISA plate (Phospho(Thr231)/total tau kit, K151221D, Meso Scale Discovery) according to the manufacturer’s protocol. Comparing total tau quantifications by WB and ELISA gave insights into the amount of tau oligomers in each sample.

### Proteinase K digestion

SARK and HMW tau seeds (2 µg total Tau per condition) were incubated with 0-10 µg/ml proteinase K (AM246, ThermoFisher) in 10 mM Tris-HCl, pH 7.4, for 10 min at RT as described previously [37]. Digestion was stopped by denaturing samples 5 min at 95°C as described above for WB. Digested samples were loaded onto a NuPAGE 4-12% Tris/Bis gel (ThermoFisher) with NuPAGE MES running buffer (ThermoFisher). Anti-total tau antibody (1: 5000, A0024, DAKO) was used as primary antibody.

### Negative staining electron microscopy

For electron microscopy (EM), 0.3 µg monomeric equivalent of total tau was loaded onto F/C 300 mesh Nickel grids for two minutes before excess solution was wicked away. After a quick wash with ultrapure water, 3 µl of 2% uranyl acetate was applied to the grid and incubated for 90 seconds, the excess wicked away, and grids were air-dried before imaging. Images were taken on a ThermoFisher Scientific T12 electron microscope at the Harvard Medical School Molecular Electron Microscopy Suite (HMS MEMS) at magnifications of 67,000x and 150,000x.

For quantification of fibril-like structures, a JEOL 1011 electron microscope was used to acquire 20 systematically randomly sampled grid squares at magnification 100,000x where the microscopist was blinded to sample identities. Fibrils were characterized and selected using PHF-resembling criteria: 10-20 nm in width and at least half a turn of the helical filament (> ∼35 nm) in length. Quantification data has been reproduced by two distinct experimenters on four different AD cases, with consistent results.

### *In vitro* seeding assay

For *in vitro* seeding assay, we used the widely used FRET-biosensor assay [17, 24] with cells stably expressing the PS19-mutant tau repeat domain conjugated to either cyan fluorescent protein (CFP) or yellow fluorescent protein (YFP) (TauRD-P301S-CFP/YFP). Briefly, cells were plated on 96-well plates (Costar, previously coated with 1:20 poly-D-lysine) at a density of 40,000 cells per well and cultured for 24 hours at 37°C, 5% CO_2_ in Dulbecco’s Modified Eagle Medium (DMEM), 10% fetal bovine serum (FBS) and 1% penicillin-streptomycin. SARK and HMW tau extracts (8 ng total tau per well) were incubated with 1% lipofectamine 2000 transfection reagent (ThermoFisher) in opti-MEM (ThermoFisher, final volume of 50 µl per well) for 20 min at RT before being added to the cells. After 24 hours, cells were collected for subsequent flow cytometer analysis of seeding i.e. FRET signal. Medium was removed, 50 µl trypsin 1X was added to each well for 5 min at 37°C, and the reaction was stopped by adding 150 µl fresh medium. Cells were transferred to 96-well U-bottom plates (Corning), pelleted at 1500 rpm for 10 min, resuspended in 2% paraformaldehyde (PFA) in PBS for 20 min at RT and pelleted again at 1500 rpm for 10 min. Cells were finally resuspended in 150 µl per well PBS, and run and analyzed on the MACSQuant VYB (Miltenyi) flow cytometer as previously described [17]. For each well, tau seeding value was calculated by multiplying the percentage of FRET-positive cells by the median fluorescence intensity of that FRET-positive population (integrated FRET density or IFD). Each sample was loaded in triplicate and 3 experiments were performed independently. Whether tau samples were sonicated or not is specified in the legend of each figure.

### *In vivo* seeding in tau transgenic mice

Three-month-old male PS19 tau transgenic mice (B6;C3-Tg(Prnp-MAPT*P301S)PS19Vle/J, Jackson Laboratory) overexpressing the human 1N4R isoform of tau with a P301S mutation were used. Following anesthesia with 1.5% (vol/vol) isoflurane mice were fixed in a stereotaxic frame, ophthalmic ointment was placed on the eyes, and lidocaine hydrochloride (2 mg/kg) was subcutaneously administered under the scalp skin before the beginning of surgery. Mice were intracerebrally injected into CA1 of the hippocampus (anteroposterior -2.4 mm from bregma, mediolateral +/- 1.5 mm from midline and dorsoventral -1.6 mm from the skull surface) using a 33-gauge blunt-tip needle fixed to a Hamilton syringe. Each mouse was bilaterally injected with 1µg non-sonicated SARK (n=8: 3 with #1892, 5 with #2399) or HMW (n=8: 3 with #1892, 5 with #2399) tau diluted in PBS (2.33 µl/site) at a flow rate of 0.2 µl/min. SARK and HMW samples immunodepleted for total tau (n=2/group), as well as PBS alone (n=7) were used as controls. After surgery, the skin was sutured, and buprenorphine hydrocholoride (0.05 mg/ml) subcutaneously administered every 12 hours for 72 hours. Acetaminophen (300 mg/kg) was added to drinking water *ad libitum* for 72 hours.

Three months after injection mice were euthanized with CO_2_ and transcardially perfused with ice-cold PBS for 5 min at 20 ml/min. The brains were removed, and the hemispheres separated. The left hemisphere was post-fixed for 24 hours in 4% PFA and cryoprotected in 30% sucrose for 72 hours before sectioning. 40 µm-thick coronal sections were sliced using a freezing microtome and were collected in series at 400 µm intervals.

Experiments were also performed in Tau22 mice, which express human tau with a double mutation G272V/P301S under a neuron-specific Thy1.2 promoter. Because of the more aggressive phenotype of this line, Tau22 mice of either sex were injected at 2 months of age, and were euthanized 2 months after injection and processed similarly to injected PS19 mice (n=3/group). SARK samples were hand-sonicated 60 pulses at power 2 prior to injection into Tau22 mice to assess the effect of sonication *in vivo*.

All animal care, housing and experiments were performed in compliance with guidelines established by the Massachusetts General Hospital institutional animal care and use committee and in accordance with the National Institutes of Health Guide for the Care and Use of Laboratory Animals.

### Immunohistochemistry

Free-floating 40 µm-thick coronal sections were washed in 0.2% Triton X-100 PBS and incubated for 30 min in 0.3% hydrogen peroxide. After washing in 0.2% Triton X-100 PBS, sections were blocked in 4.5% normal goat serum (Vector Laboratories) in 0.2% Triton X-100 PBS for 1 hour then incubated in primary antibody solution (Table S1) in blocking solution at 4°C overnight. The next day, sections were washed in 0.2% Triton X-100 PBS and incubated in secondary antibody solution for 1 hour before amplification of the signal by incubation in Vectastain ABC kit (1:400, Vector Laboratories) for 1 hour. After wash in 0.2% Triton X-100 PBS, sections were incubated for 1 min in 3,3’-diaminobenzidine (DAB, Vector Laboratories), then mounted on Superfrost Plus slides (Fisher Scientific), dehydrated in increasing concentrations of ethanol/xylene and coverslipped with Eukitt (Sigma-Aldrich).

Human formalin-fixed paraffin-embedded brain tissue sections from the contralateral hemisphere of AD cases #1892 and #2399 were stained for microglia (Iba1, abcam, 1:500) using DAB and counterstained with hematoxylin following the Leica autostainer protocol (Bond RX autostainer, Leica Biosystems). Slides were dehydrated and coverslipped with Permount mounting medium (ThermoFisher).

### Immunofluorescence

Free-floating 40 µm-thick coronal sections were washed in PBS, blocked with 4.5% normal goat serum in 0.2% Triton X-100 PBS for 1 hour then incubated in primary antibody solution at 4°C overnight. The next day, sections were washed in PBS and incubated in secondary antibody solution for 1 hour before being mounted on Superfrost Plus slides and coverslipped with DAPI-containing Fluoromount-G (0100-20, Southern Biotech).

### Thioflavin-S staining

Free-floating 40 µm-thick coronal sections were washed in PBS, mounted on Superfrost Plus slides and let dry overnight. The following day, sections were quickly rinsed in PBS 1X before incubation in 0.05% Thioflavin-S (ThioS, T1892-25G, Sigma-Aldrich) in 50% ethanol for 8 min in the dark. Sections were then briefly rinsed in 80% ethanol and transferred to dH_2_O for 5 minutes prior to coverslipping.

### Image analysis

Immunohistology images were acquired at 20x using a NanoZoomer (Hamamatsu). Manual segmentation of the different brain regions and thresholding of each channel was then performed on 3 sections adjacent to the injection site using QuPath software. Staining-positive cells were manually counted using QuPath cell counter plugin. Quantification in the perirhinal/entorhinal cortex was performed on 6 sections spanning the dorsal and ventral hippocampus. All quantifications were performed blinded to the injection groups. Glial representative images were acquired on a FV3000 confocal microscope (Olympus).

### Super resolution microscopy and quantification of synaptic densities

Tissue sections prepared for super resolution imaging were mounted on charged slides as described and coverslipped with ProLong Glass Antifade mounting medium (P36980, Invitrogen) using 170 ± 5 µm No. 1.5H High Precision cover glasses. Bassoon- and PSD95-immunopositive puncta were captured using the Zeiss Elyra 7 super resolution microscope with Lattice SIM2. Images were acquired using a 63x oil objective. 3.91 µm z-stacks were imaged at a step-size of 0.126 µm, with laser power 0.9% and 90 ms exposure, and subsequently processed using the SIM2 ‘Low contrast’ reconstruction settings. Total number of Bassoon- and PSD95-puncta per z-stack were determined by thresholding using Imaris Software and normalized per mm^3^ of tissue.

### Statistical analysis

Statistical analyses were performed using GraphPad Prism software. The normality of the distribution and homoscedasticity were checked prior to analysis and the corresponding parametric or non-parametric statistical tests performed. Paired tests were used for the comparison between SARK and HMW tau derived from the same AD cases. For *in vivo* experiments, statistical tests were followed by post-hoc multiple comparisons between all three injection groups to account for both the effect of the injected tau seeds (HMW tau vs PBS, SARK tau vs PBS) and their nature (HMW tau vs SARK tau). Data are represented as mean and standard error of mean (mean±SEM). Outliers identified by GraphPad Prism software ‘Identify outlier’ plug-in (ROUT Q = 5%) were removed from the analysis.

## Results

### AD-derived SARK and HMW tau have different tau compositions

To evaluate tau species’ heterogeneity within the brain of AD patients, we extracted tau from the cortex (BA 8/9 or BA 7, 5 g) of 8 AD cases using two different methods. Cases were chosen based on the presence of Braak stage V-VI, and their high seeding competency as previously described [17] (Table 1). Dorsolateral prefrontal cortex (Brodmann areas 8 and 9) or parietal lobe (Brodmann area 7) were utilized as available in each case, as both association areas are strongly affected by tau pathology in advanced AD. For each AD case we isolated both i) SARK tau corresponding to more fibrillar species of tau using the well characterized sarkosyl-insoluble extraction method [22, 42], and ii) HMW tau corresponding to tau oligomers using size-exclusion chromatographyon PBS soluble tissue fractions [17, 57] (Fig. 1a). Quantification of total tau load by both denaturing WB and non-denaturing ELISA reveals a significant difference between SARK and HMW tau preparations (Supplementary Fig. 1a-b) demonstrating that there is more fibrillar tau per tissue weight than there is soluble HMW species. Interestingly, the ratio of tau as measured by WB and ELISA which reflects tau oligomerization is significantly higher in the HMW samples compared to SARK samples (Supplementary Fig. 1c) confirming the distinct folding of these two species, as reflected by antibody access in an ELISA assay. To further evaluate the structural characteristics of these tau species, we incubated SARK and HMW tau from one AD case in increasing concentrations of proteinase K, a broad-spectrum serine protease, and looked at the digestion profiles by total tau WB. As expected, a higher concentration of proteinase K is needed to digest SARK tau compared to HMW tau (Fig. 1b-c). These data are consistent with electron microscopy characterization, where HMW tau displays amorphous material with few short fibrillar species suggesting mainly oligomeric forms of tau, while SARK tau presents a high proportion of short and long twisted tau fibrils characteristics of paired-helical filaments (PHFs) (Fig. 1d). Because SARK tau samples are usually used sonicated to enhance their bioactivity [22, 42], we evaluated the influence of sonication on both SARK and HMW tau species (Supplementary Fig. 1d). While fibrillar structures are absent from the sonicated HMW tau sample and unchanged in the sonicated SARK tau sample, quantification of PHF-like structures on negative stain EM images confirms the very low abundance of fibrillar tau in the non-sonicated HMW sample compared to non-sonicated SARK sample despite comparable fibril sizes (Supplementary Fig. 1e-f). These observations strengthen the idea of multiple tau species in the AD brain with SARK tau containing mostly fibrillar and HMW tau essentially consisting of oligomeric forms of tau.

**Fig. 1:**
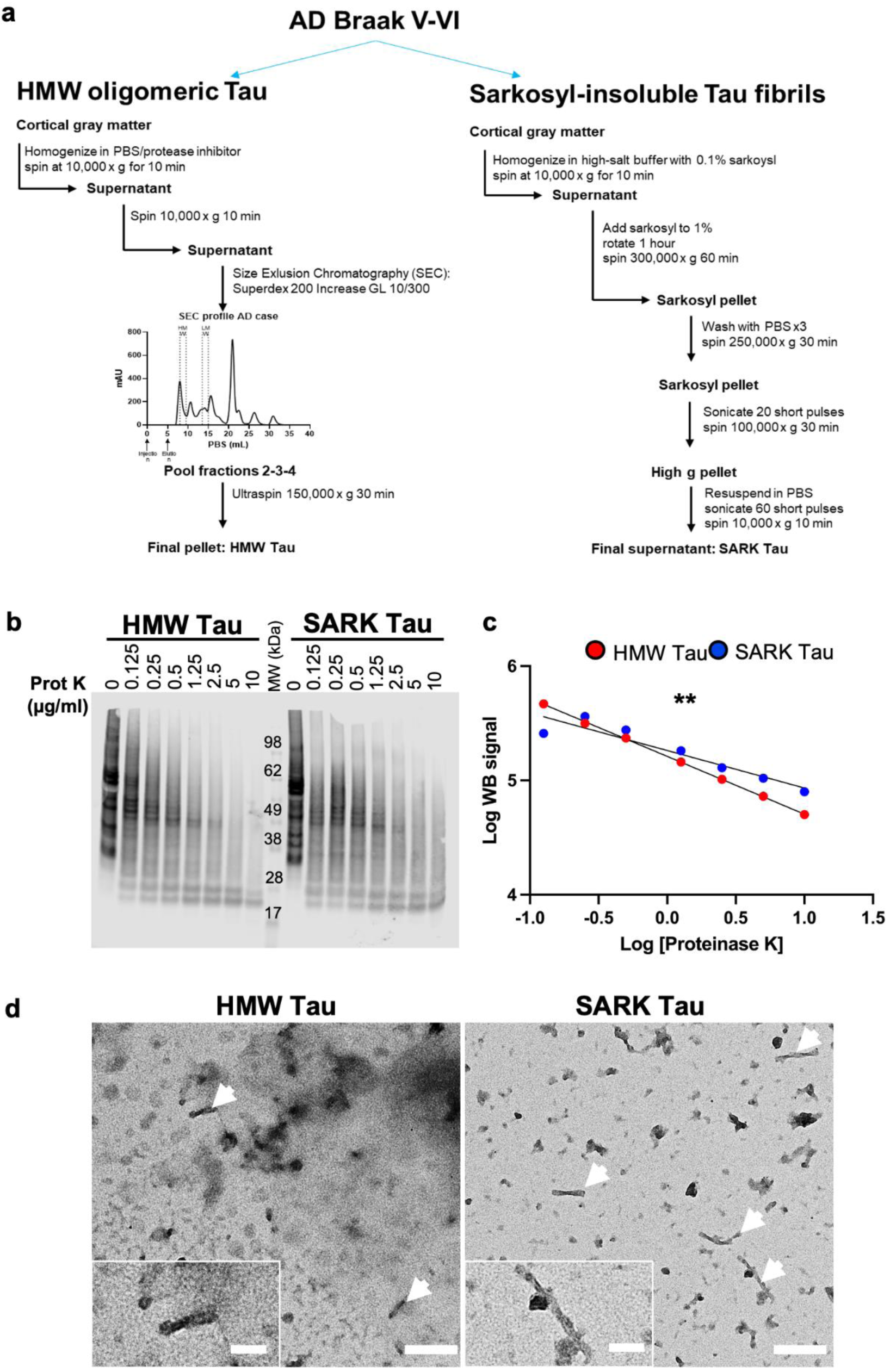
SARK tau is mainly fibrillar while HMW tau is mainly oligomeric. **a.** Schematic of the experimental workflow to extract tau oligomers [17] (HMW Tau) and tau fibrils [42] (SARK tau) from the cortical gray matter of 8 AD cases of Braak V-VI. **b.** Total tau WB (DAKO) of SARK and HMW tau samples after incubation with increasing concentrations of proteinase K. **c.** Quantification of the WB signal intensity after proteinase K treatment. Log-transformed data are plotted, and the slopes compared. Linear regression testing for the equality of slopes, **p<0.01. **d.** Representative negative stain EM images of non-sonicated samples from AD case #2399 used for *in vivo* injections. HMW tau samples present mainly as amorphous material and a few PHFs (white arrows) while PHFs are more abundant in the SARK tau samples. Scale bar = 200 nm and 100 nm for inserts. MW = molecular weight.

**Table 1:** AD cases demographics and characteristics used in this study. SARK and HMW tau extracted from AD cases #1892 and #2399 were used for injections into PS19 mice. NPDX = neuropathological diagnostic, ADNC = AD neuropathological changes, CVD = cerebrovascular disease, LBD = Lewy body disease, CAA – cerebral amyloid angiopathy.

| #ADRC | Brain region | Age at death | Sex | NPDX1 | NPDX2 | NPDX3 | Braak Stage | Thal Stage | Note |
| --- | --- | --- | --- | --- | --- | --- | --- | --- | --- |
| 1648 | BA 8/9 | 75 | Female | ADNC | CVD |  | VI | 4 |  |
| 1853 | BA 8/9 | 83 | Female | ADNC | LBD | CVD | VI | 5 |  |
| 1892 | BA 8/9 | 67 | Male | ADNC |  |  | VI | 5 | Injected |
| 1971 | BA 8/9 | 76 | Male | ADNC | CVD | CAA | VI | 4 |  |
| 2014 | BA 8/9 | 84 | Male | ADNC | CAA |  | VI | 5 |  |
| 2157 | BA 8/9 | 63 | Female | ADNC | Arteriolosclerosis |  | VI | 5 |  |
| 2399 | BA 7 | 74 | Male | ADNC | CVD |  | V | 5 | injected |
| 2520 | BA 7 | 61 | Female | ADNC | CVD | CAA | VI | 5 |  |

### AD-derived SARK and HMW tau have similar seeding activities *in vitro*

To biologically compare SARK and HMW tau, we analyzed their bioactivity by performing a seeding assay using the well-established FRET-biosensor cell line [24]. SARK and HMW tau seeds were lipofectamine-transfected into cells expressing a tau-biosensor construct and incubated for 24 hours before flow cytometry analysis of the FRET signal (Fig. 2a). For each condition, 8 ng monomeric equivalent of tau was added per well. Data show that SARK and HMW tau have similar seeding activities in this specific assay whether SARK tau was sonicated or not (Fig. 2b), and this is consistent among our 8 AD cases (average seeding ratio between HMW and non-sonicated SARK tau samples = 1.16 ; average seeding ratio between HMW and sonicated SARK tau samples = 1.05). While sonication affects the number of PHF-like structures in the HMW tau sample (Supplementary Fig. 1d), it does not affect its seeding capacity (Supplementary Fig. 1g) suggesting that fibrillar structures are not driving the bioactivity of the HMW tau sample, and thus strengthening the difference between SARK and HMW tau samples.

**Fig. 2:**
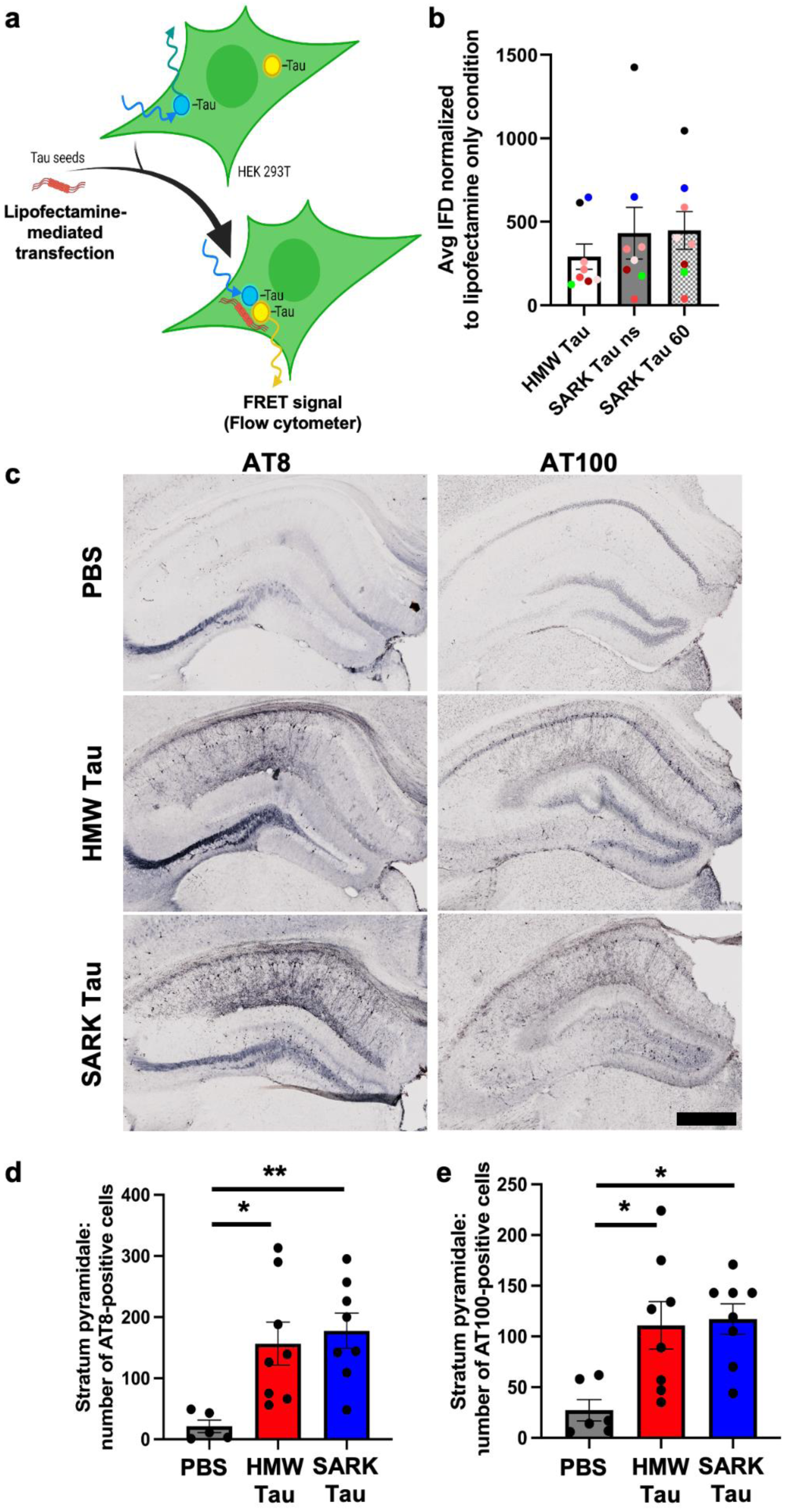
Tau oligomers and tau fibrils have similar bioactivities *in vitro* and *in vivo*. **a.** Schematic of the FRET-biosensor seeding assay. HEK293T cells stably express the tau repeat domain fused to either CFP or YFP. When transfected with bioactive tau seeds, these constructs aggregate and become close enough to generate a FRET signal quantifiable by flow cytometry. Here, SARK and HMW tau samples from 8 AD cases were normalized to total tau amounts (8 ng monomer equivalent per well) before being added to the cells. Artwork was created using BioRender. **b.** Quantification of the seeding activity by flow cytometry 24 hours after transfection. Integrated FRET densities were normalized to the lipofectamine only negative control. Data represented as mean±SEM from 3 independent experiments, one color per case, Wilcoxon matched-pairs test, non-significant. Cases further used for in vivo experiments have been highlighted in green for #1892 and blue for #2399. ns=non-sonicated ; 60=sonicated 60 pulses. **c.** Representative images of phosphorylated AT8 and AT100 tau pathology in the dorsal hippocampus of injected PS19 mice 3 months after injection. Scale bar = 500 µm. **d.** Quantification of the number of AT8-positive cells in the pyramidal layer (CA pyramidale) of the dorsal hippocampus of PS19 mice 3 months after injection. **e.** Quantification of the number of AT100-positive cells in the pyramidal layer (CA pyramidale) of the dorsal hippocampus of PS19 mice 3 months after injection. Data represented as mean±SEM, Kruskal-Wallis, Dunn’s multiple comparison, *p<0.05.

### Tau pathology is enhanced after the injection of AD-derived SARK and HMW tau seeds *in vivo*

We next wondered how a complex system like a mouse brain would impact the bioactivity of SARK and HMW tau. We intracerebrally injected the different tau species isolated from two AD cases into the dorsal hippocampus of male PS19 Tau transgenic mice. To assess if there were major differences between tau that was isolated from different AD brains, the experimental design included injecting SARK or HMW tau isolated from AD case #1892 (3 animals) or AD case #2399 (5 animals). Both cases displayed similar seeding activity *in vitro*. PS19 mice express the P301S-mutated human 1N4R tau isoform under the Prnp prion promoter resulting in progressive tau pathology with the appearance of NFTs by 6 months of age [66]. By injecting PS19 mice at 3 months of age and sacrificing them 3 months later, we anticipated a faster development of tau pathology and used it as readout for seeding activity. PBS-injected mice reflect spontaneous tau pathology linked to the expression of the mutant human tau transgene. As expected, in 6-month-old PS19 mice, PBS-injected mice display sparse hyperphosphorylated AT8- and fewer AT100-positive neurons in the pyramidal layer of the dorsal hippocampus around the injection site. Three months after the injection of SARK or HMW tau, PS19 mice exhibited dense hippocampal AT8- and AT100-tau-positive staining in the vicinity of the injection site (Fig. 2c). Quantification of the number of AT8- and AT100-positive cells in the pyramidal layer of the dorsal hippocampus confirms that both SARK and HMW tau samples promote neuronal tau pathology in PS19 mice by about 8-fold compared to PBS-injected animals (Fig. 2d-e). Thus, SARK and HMW tau seeds derived from the same AD brain trigger AT8 and AT100 tau pathology to the same extent locally at the injection site suggesting both these tau species have similar local seeding activities.

We next assessed the regional distribution of tau pathology in the different injection groups which could reflect the behavior of the injected tau seeds and their respective consequences on endogenous tau. Quantification of the percentage of AT8-positive area in the *stratum oriens* and *stratum radiatum* in the dorsal hippocampus reflects the axonal and dendritic tau pathologies close to the injection site (Supplementary Fig. 2a). Interestingly, while both SARK and HMW tau increase the number of AT8-positive hippocampal neurons compared to baseline, the SARK tau injected mice display statistically increased dendritic tau pathology compared to PBS-injected PS19 (Fig. 2c and Supplementary Fig. 2b-c) suggesting a differential intracellular distribution of tau pathology after seeding by HMW or SARK tau.

### Tau pathology propagation to distal, anatomically connected brain areas occurs 3 months after AD-derived SARK and HMW tau injection

In human AD cases, NFT involvement is not confined to a specific brain region, but slowly progresses throughout the brain in a hierarchical pattern referred to as Braak stages I-VI according to the brain region affected by AT8-positive NFTs [9]. This observation gave rise to the idea that pathological tau protein(s) propagate along connected brain regions and trigger the formation of tau pathology in recipient cells. We here showed that SARK and HMW tau species can both trigger tau pathology in hippocampal recipient cells surrounding the injection site. We next evaluated whether tau pathology is increased in brain regions distal to the injection site and use it as a readout for tau spreading. We used the AT8 antibody, as it is commonly used in neuropathology to assess Braak staging in AD brains, and we quantified the number of AT8-positive cells in the overlying isocortex including the retrosplenial, motor and somatosensory cortices, and in the combined perirhinal and entorhinal cortices along the hippocampal formation. Because the injection needle goes through the isocortex during intrahippocampal injection, tau pathology in this region reflects either the diffusion of the tau seeds during injection, or local projections. On the other hand, the distal peri-/entorhinal cortices are synaptically connected to the hippocampus, so the extent of tau pathology in this brain region is an indicator of tau seeds’ spreading along anatomical pathways. Three months after injection, PBS-injected PS19 mice present very few AT8-positive cells in both the overlying and entorhinal cortices. Interestingly, 3 months after intrahippocampal injection of HMW tau AT8 tau pathology seems to increase in both brain regions while after SARK tau injection AT8 tau pathology is present only in the overlying isocortex (Fig. 3a). Quantification of the number of AT8- and pS422-positive cells in the isocortex confirms the almost 10-fold and 2-fold increase, respectively, in SARK and HMW tau-injected animals compared to the PBS-injected PS19 and shows no statistical difference between the two tau-injected groups (Fig. 3b). Although the number of neurons in the entorhinal/perirhinal cortex area had a fair amount of animal-to-animal variability, in some cases the number of AT8- and pS422-positive cells in the peri- /entorhinal of the HMW tau-injected group 3 months after injection were noticeably higher than PBS or even SARK injected animals, although a difference between the latter did not reach statistical significance (Fig. 3c, e). In accordance with these data, the number of ThioS-positive cells in the entorhinal cortex after HMW tau injection was significantly higher than the ‘endogenous’ levels reflected by PBS injection (see Fig. 4d). These results suggest that propagation and/or seeding kinetics in distal brain regions occur after both SARK and HMW tau seeds.

**Fig. 3:**
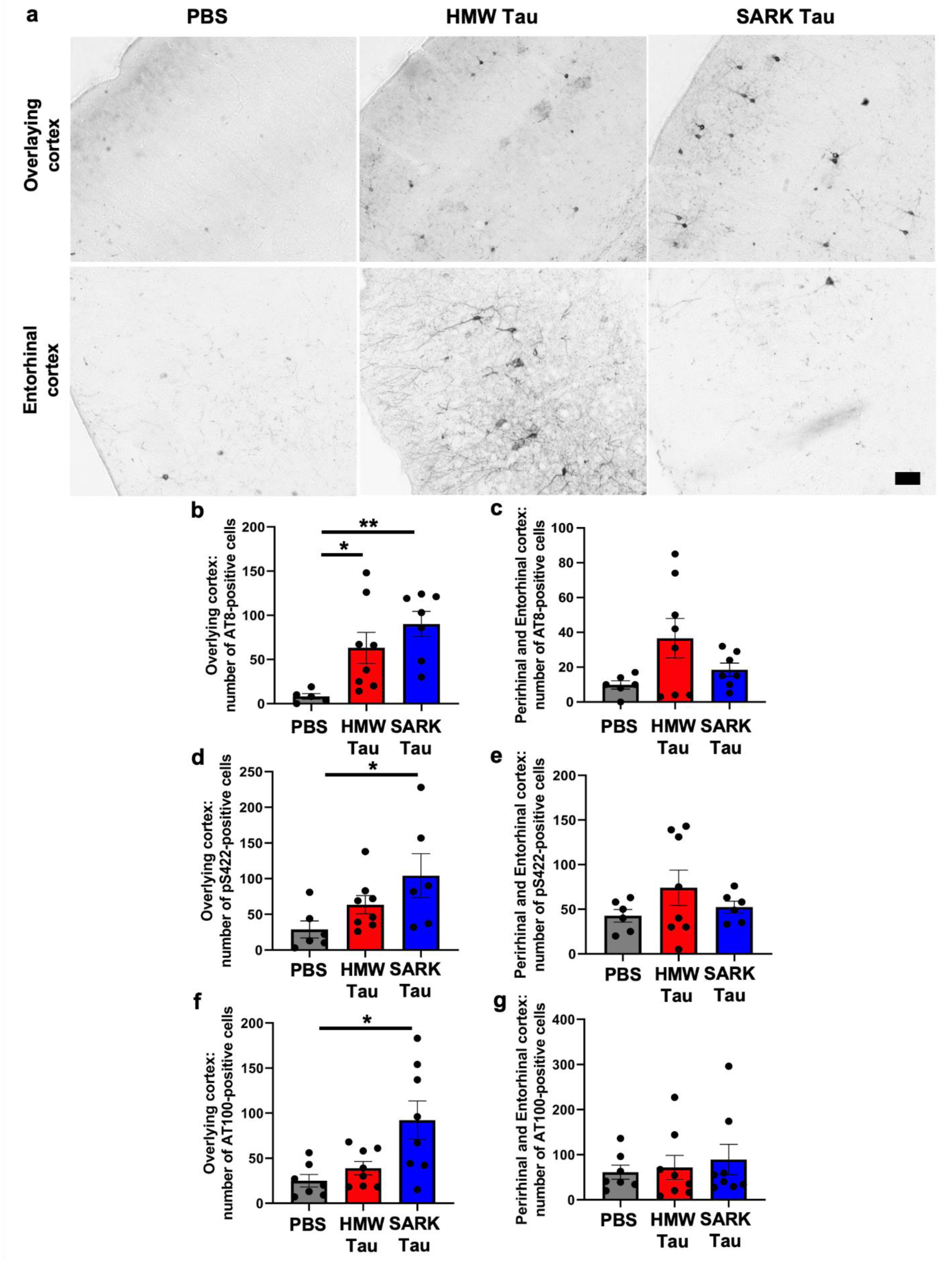
Tau oligomers and tau fibrils both spread to interconnected distal brain regions. **a.** Representative images of phosphorylated AT8 tau pathology in the overlying isocortex (upper panel) and entorhinal cortex (lower panel) of PS19 mice 3 months after injection. Scale bar = 50 µm. **b-c.** Quantification of the number of AT8-positive cells in the cortical areas overlying the dorsal hippocampus and in the perirhinal/entorhinal cortices, respectively, in PS19 mice 3 months after injection. The overlying isocortex comprises the retrosplenial, motor and somatosensory cortices. **d-e.** Quantification of the number of pS422-positive cells in the cortical areas overlying the dorsal hippocampus and in the perirhinal/entorhinal cortices, respectively, in PS19 mice 3 months after injection. **f-g.** Quantification of the number of AT100-positive cells in the cortical areas overlying the dorsal hippocampus and in the perirhinal/entorhinal cortices, respectively, in PS19 mice 3 months after injection. Data represented as mean±SEM, Kruskal-Wallis, Dunn’s multiple comparison, *p<0.05, **p<0.01.

**Fig. 4:**
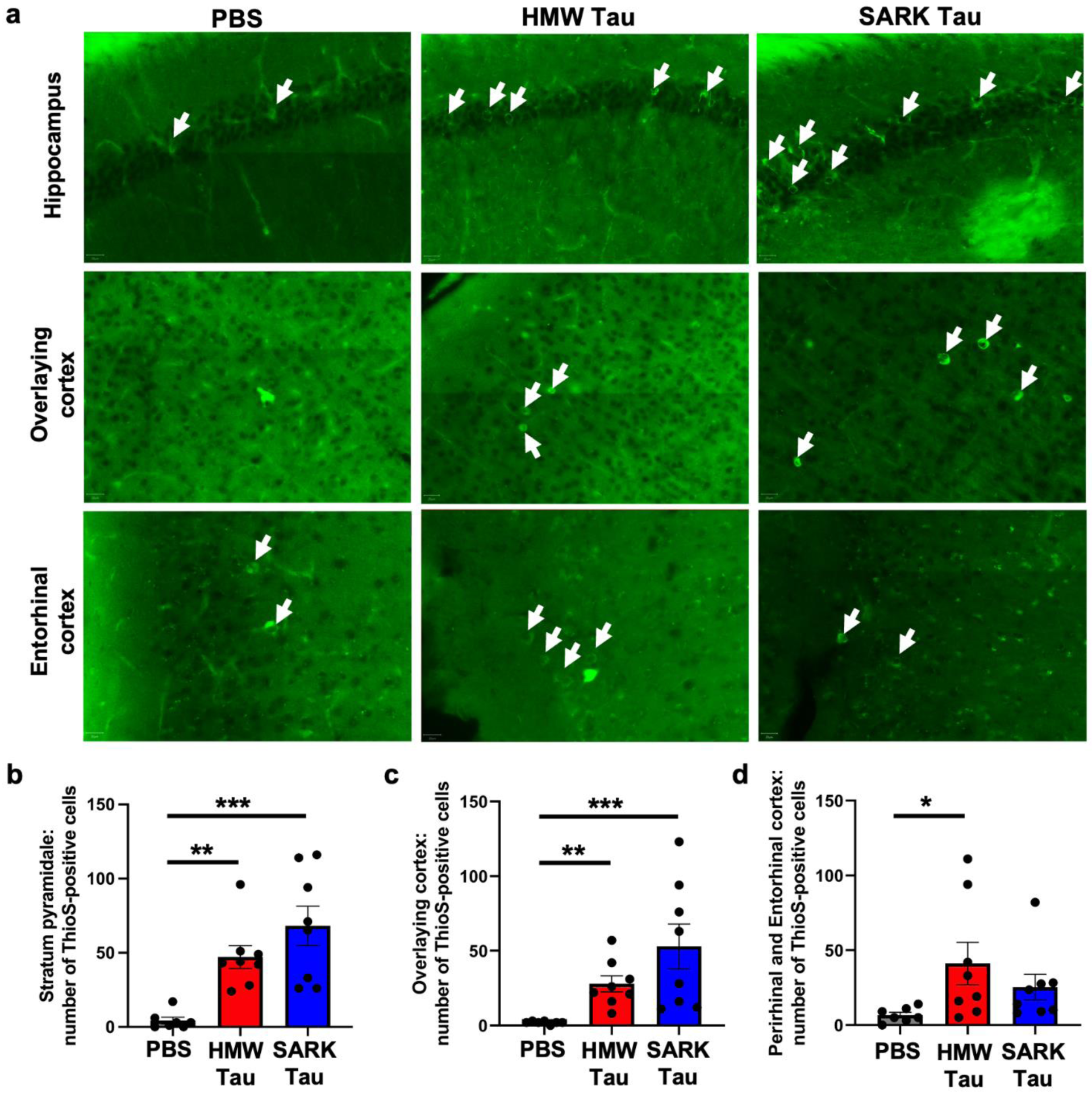
Tau oligomers and tau fibrils accelerate tau pathology maturation. **a.** Representative images of ThioS-positive staining (green) in the dorsal hippocampus, iso- and entorhinal cortices in PS19 mice 3 months after injection. White arrows show ThioS-positive neurons. Scale bars = 20 µm. **b-d.** Quantification of the number of ThioS-positive cells in the pyramidal layer of the dorsal hippocampus, the isocortex and the peri-/entorhinal cortices respectively in PS19 mice 3 months after injection. Data represented as mean±SEM, Kruskal-Wallis, Dunn’s multiple comparison, *p<0.05, **p<0.01, ***p<0.001.

We further examined the propagation and seeding abilities of the SARK and HMW tau seeds by evaluating the maturation of tau pathology in the different injection groups. We thus investigated protein aggregation using Thioflavin-S (ThioS), a β-pleated sheet conformation-specific fluorescent dye, in our 3 different regions of interest (Fig. 4a). As expected, ThioS-positive cells are almost non-existent in the PBS-injected animals, but abundantly present in the pyramidal layer of the hippocampus and the isocortex in both SARK and HMW tau-injected PS19 mice 3 months after injection (Fig. 4b-c). These data suggest that the presence of SARK and HMW tau seeds accelerate the maturation of tau pathology, thus being involved in the progression of the disease. Consistent with our results of AT8 tau pathology in the peri-/ento-rhinal, the number of ThioS-positive cells is increased only in the HMW tau-injected animals compared to the PBS group (Fig. 4d) confirming that SARK and HMW tau seeds have different spreading properties to synaptically connected brain regions with HMW tau-induced pathology progressing faster.

Mature tau aggregates or NFTs are commonly considered as the endpoint of pathological tau maturation [38] and appear to be highly phosphorylated at multiple sites [54, 60]. At the time of sacrifice (3 months after injection, so 6 months of age) PS19 mice already exhibit endogenous tau pathology. We thus evaluated the correlation between ThioS-positive tau aggregates and three tau phosphorylation epitopes including AT8, AT100 and the fibrillar-tau associated pS422 [4, 11, 23, 33, 49, 67] in our three regions of interest. Curiously, while the number of ThioS-positive cells does not correlate with the number of AT8- or AT100-positive cells in any brain region of interest (Supplementary Fig. 2d-i), it does correlate with the number of pS422-positive cells in all the three regions we looked at (Supplementary Fig. 2j-l).

### Microglial response differs according to the tau seed

To further evaluate the downstream effects of the different tau seeds, we looked at the glial response in proximity to the injection site. Tissue was stained with the commonly used GFAP marker for astrocytic reactivity and Iba1 for microglial response. While the GFAP-covered area is significantly increased in the HMW-injected animals compared to PBS-injected mice (Fig. 5a), quantification of the Iba1-covered area does not show any difference among groups (Fig. 5b).

**Fig. 5:**
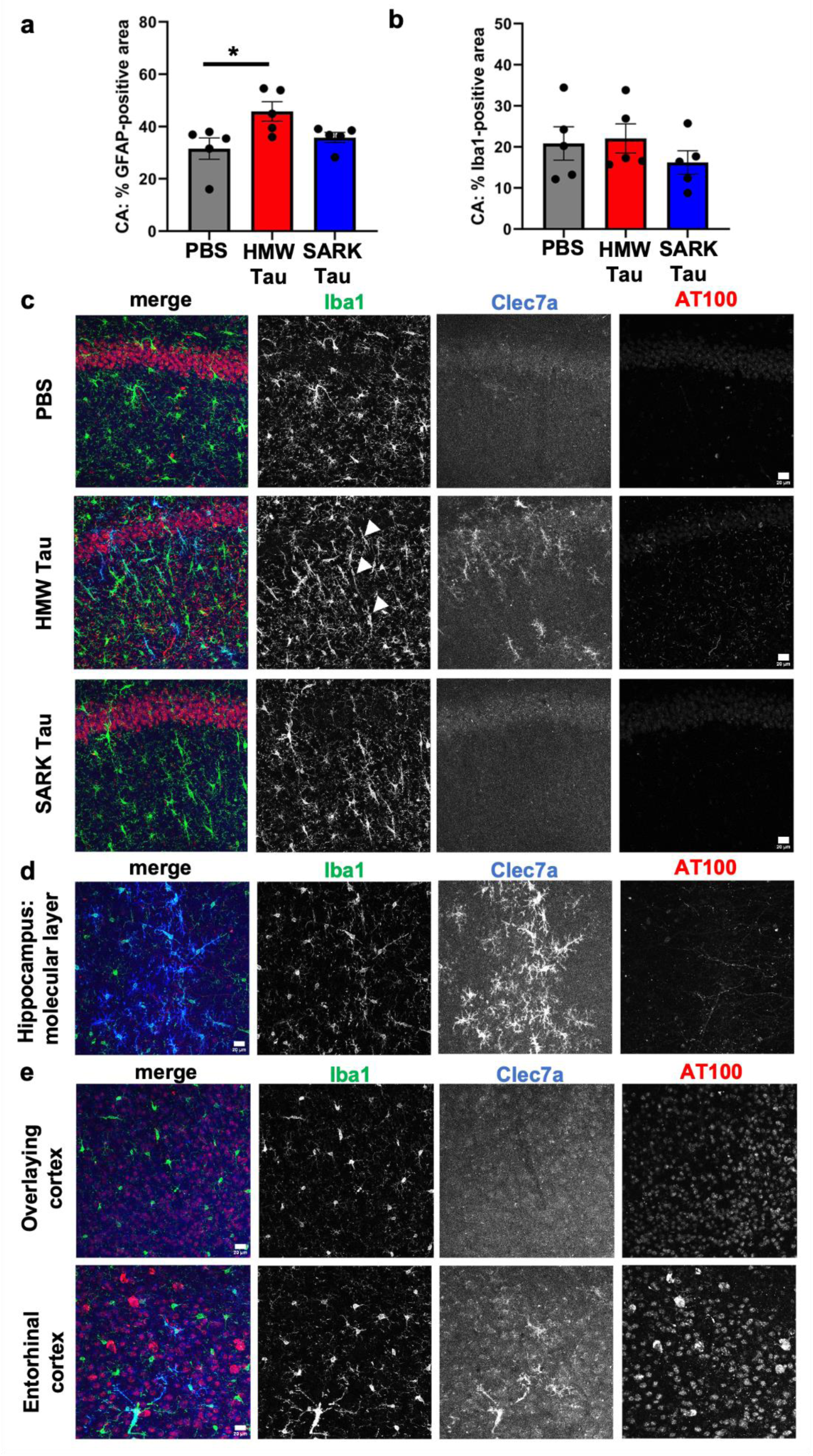
Only Tau oligomers induce rod-like and Clec7a-positive microglia after injection. **a.** Quantification of the percentage of GFAP-positive astrocyte covered area in the dorsal hippocampus of PS19 mice 3 months after injection. **b.** Quantification of the percentage of Iba1-positive microglia covered area in the dorsal hippocampus of PS19 mice 3 months after injection. **c.** Representative images of Clec7a-positive (blue) and Iba1-positive (green) rod-shaped microglia in the dorsal hippocampus of PS19 mice 3 months after injection. Rod-microglia (white arrowheads) is not necessarily along pathological AT100-positive neurons (red), and Clec7a-positive microglia are only present in the HMW-tau injected animals. **d.** Representative images of Clec7a- (blue) and Iba1- (green) double positive microglia and their localization to AT100-positive neurons (red) in the hippocampus of HMW tau-injected PS19 mice 3 months after injection. Clec7a-positive microglia are particularly abundant in the molecular layer of the hippocampus containing the axons of CA1 pyramidal neurons. **e.** Representative images of Clec7a- (blue) and Iba1- (green) double positive microglia in relation to AT100-positive cells (red) in the cortex overlying the dorsal hippocampus and in the entorhinal cortex of HMW tau-injected PS19 mice 3 months after injection. Some Clec7a-positive microglia are observed in the entorhinal cortex, direct projection site of the hippocampal neurons suggesting Clec7a-positive microglia as a readout for oligomeric HMW tau propagation. Scale bars = 20 µm. Data represented as mean±SEM, Kruskal-Wallis, Dunn’s multiple comparison, *p<0.05.

Curiously, despite similar Iba1-covered areas, HMW tau injected animals present a particular microglial phenotype reminiscent of rod-like microglia featuring an elongated soma and nucleus as well as shortened processes (Fig. 5c). Rod-like microglia have been reported in various diseases including AD, cerebral ischemia and demyelinating lesions [6, 34]. Interestingly, the two AD cases used here present Iba1-positive rod microglia in the grey matter of the parietal cortex (BA 7) from where SARK and HMW tau were extracted (Supplementary Fig. 3a-b). In the HMW tau-injected animals, rod microglia are mainly localized to the *stratum radiatum* along neuronal dendrites projecting to cortical areas. These rod microglia are not necessarily surrounding AT100-positive neurons, suggesting that neuronal tau inclusions themselves may not be the primary trigger for this microglial phenotype.

We next performed additional immunostaining to better define rod microglia. Based on RNAseq data from previous work [31, 47], we selected Clec7a (Dectin-1) as a marker of disease-associated microglia. Clec7a is a C-type lectin receptor expressed by myeloid cells including microglia, and is involved in antifungal responses, phagocytosis and pro-inflammatory response [40, 59]. Surprisingly, Clec7a- and Iba1-double-positive microglia are observed only in the hippocampus of HMW tau injected mice (Fig. 5c) and mainly close to the injection site. Most importantly, many of the Clec7a-positive microglia are also rod-shaped. In contrast, not all the Iba1-positive rod microglia are Clec7a-positive suggesting cellular heterogeneity within this rod-like population. Interestingly, Clec7a-positive microglia are mainly localized to the hippocampus, especially the *stratum radiatum* and the molecular layer, which contain dendrites and axons of CA1 pyramidal neurons projecting to the piriform cortex, respectively (Fig. 5d). No rod- or Clec7a-positive microglia are observed in the isocortex, but a few Clec7a-positive microglia are spotted in the entorhinal cortex (Fig. 5e) after HMW, but not SARK or PBS injections. The entorhinal cortex is a direct projection site of hippocampal neurons, suggesting that Clec7a-positive microglia may be an early readout for oligomeric HMW tau propagation.

Both rod and Clec7a-positive microglia have been hypothesized to be associated with axonal dysfunction [18, 51]. To evaluate the possibility that the appearance of rod microglia is associated with differential cellular damage after HMW but not SARK injection, we evaluated hippocampal atrophy in the different injection groups. Data show that neither the volume of the pyramidal layer nor the volume of the CA fields differs among the injection groups (Fig. 6a-b). We further quantified pre- and post-synaptic densities in the *stratum radiatum* where most of the rod- and Clec7a-positive microglia are present. Interestingly none of the Bassoon-positive pre-synaptic or PSD95-positive post-synaptic densities differ among any groups (Fig. 6c-d) suggesting that the presence of SARK and HMW tau seeds does not aggravate synapse loss in PS19 mice 3 months after injection.

**Fig. 6:**
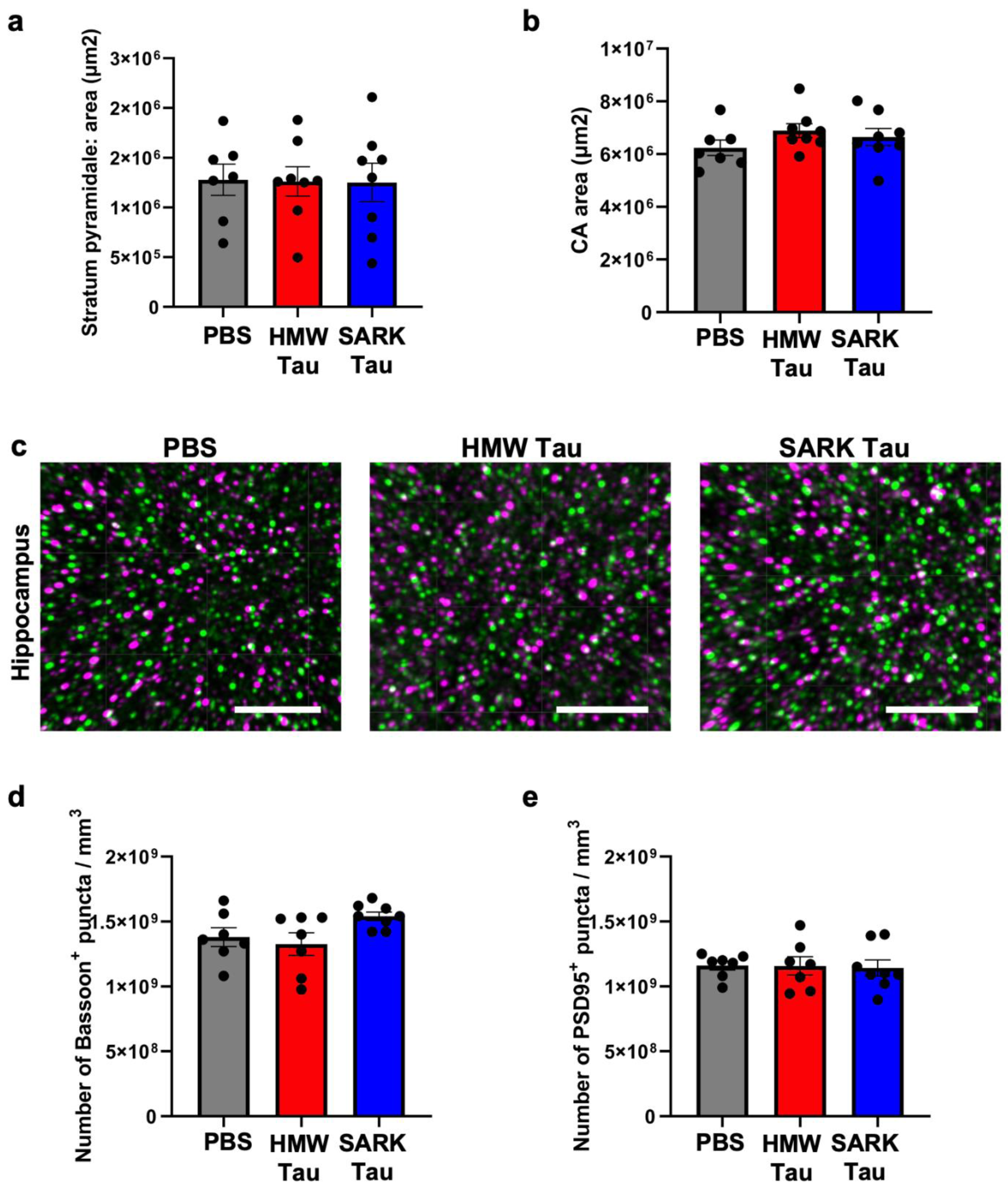
Tau oligomers and tau fibrils do not trigger evident neurodegeneration. **a.** Quantification of the area of the DAPI-positive pyramidal layer in the dorsal hippocampus of PS19 mice 3 months after injection. **b.** Quantification of the CA area of the dorsal hippocampus of PS19 mice 3 months after injection. **c.** Representative images of Bassoon-positive pre-synapses (green) and PSD95-positive post-synapses (magenta) in the *stratum radiatum* of PS19 mice 3 months after injection. Scale bar = 4 µm. **d.** Quantification of Bassoon-positive pre-synaptic density in the *stratum radiatum* of PS19 mice 3 months after injection. **e.** Quantification of PSD95-positive post-synaptic density in the *stratum radiatum* of PS19 mice 3 months after injection. Data represented as mean±SEM, Kruskal-Wallis, non-significant for a-b. Data represented as mean±SEM, One-way ANOVA, non-significant for c-d.

Taken together, these data suggest that the observed biological effects on glia following SARK and HMW tau injections do not depend on the triggered tau pathology nor cellular loss but rather reflect the intrinsic properties of the injected tau seeds.

### The effects of AD-derived SARK and HMW preparations are driven by tau

To verify the tau specificity of the effects of injecting human brain derived preparations into the mouse, we immunodepleted Tau from both SARK and HMW samples using the HT7 total human Tau antibody before running them on the biosensor seeding assay. As expected, reduction of Tau protein from both SARK and HMW samples (Supplementary Fig. 4a) almost completely abolishes *in vitro* seeding activity lowering it to the same level as SARK and HMW tau samples from a non-AD control case (Supplementary Fig. 4b-c). When immunodepleted for tau, SARK and HMW samples do not promote tau pathology in PS19 mice up to 3 months after injection (Supplementary Fig. 4d) reinforcing the tau specificity of the effect. Most importantly, only a few rod-like microglia and no Clec7a-positivity are observed in the tau immunodepleted HMW group (Supplementary Fig. 4e).

We next asked if the differential effects observed for SARK and HMW tau are intrinsic to the seeds themselves or dependent on the tau transgenic model. We thus intrahippocampally injected male and female Tau22 mice which express double-mutated G272V/P301S human tau under a neuron-specific Thy1.2 promoter [49] and where spontaneous tau pathology appears as early as 3 months of age. To evaluate *in vivo* seeding activity, we injected Tau22 mice at 2 months of age, before the appearance of spontaneous tau pathology, and we performed histological analysis 2 months after injection. As expected, SARK (here sonicated 60 short pulses) and HMW tau-injected Tau22 mice display increased AT8- and AT100-positive staining in the dorsal hippocampus compared to their PBS-injected littermates (Supplementary Fig. 5a). Interestingly, evaluation of the glial response in injected Tau22 mice reveals a similar outcome as in PS19 mice with the appearance of rod-like microglia in the HMW tau-injected animals only (Supplementary Fig. 5b). These Tau22 data complement the PS19 data by showing that tau seeds’ effect does not depend on the tau transgene that is expressed, and instead is most likely a function of the soluble or fibrillar seeds that are introduced. Additionally, similar to our *in vitro* bioactivity data, sonication does not seem to affect the *in vivo* bioactivity of the SARK tau sample. All in all, these results strengthen the idea of the co-existence of different tau seeds within an AD brain with distinct downstream effects (Fig. 7).

**Fig. 7:**
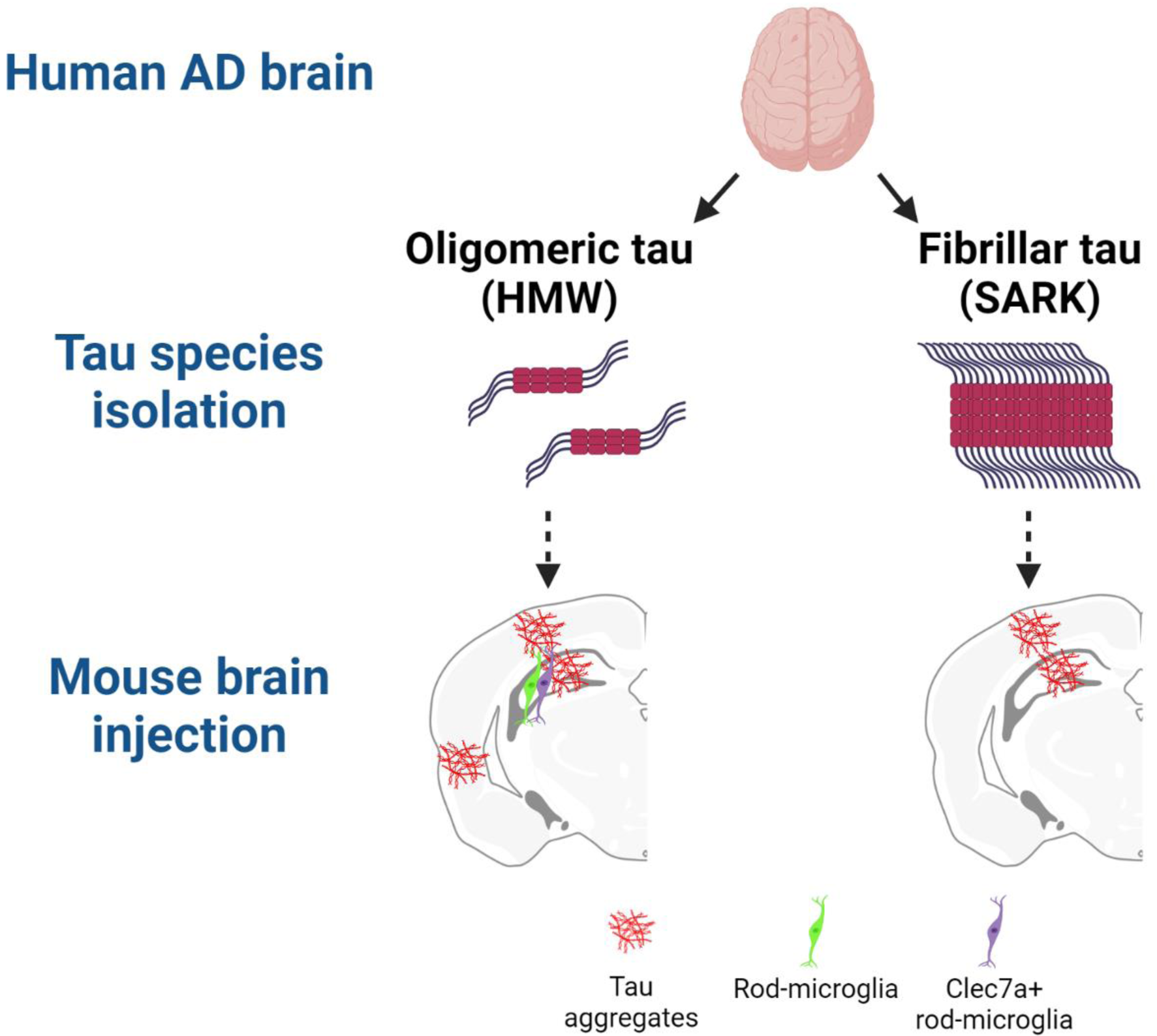
Distinct tau species induce distinct biological effects. Schematic representation of the main findings using BioRender. AD brain-derived tau species have been isolated using PBS solubility and SEC (HMW) or sarkosyl-insolubility (SARK) obtaining mostly oligomeric forms of tau and mostly fibrillar tau, respectively, after biochemical and EM characterization. Tau samples were then injected into the hippocampus of PS19 and Tau22 tau mouse models. Histological analysis of injected mouse brains revealed extensive neuronal tau pathology (AT8-, pS422-, AT100- and ThioS-positive) in the hippocampus and the overlying cortex in both injection groups, but abundant tau pathology in the peri-/entorhinal cortices as well as Clec7a-positive rod microglia at the injection site only in the HMW tau-injected animals attributing distinct biological effects to distinct tau species.

## Discussion

AD is a progressive neurodegenerative disease characterized by the presence of amyloid plaques and NFTs in the brain of cognitively impaired patients [1]. NFTs are formed by the aggregation of the microtubule-associated tau protein and their presence has been historically associated with the progression of the disease [2, 9, 19]. Tau protein is physiologically present in the brain of every individual. However, under pathological conditions tau protein undergoes various modifications leading to its accumulation and progressive oligomerization into mature aggregates [38], suggesting that multiple tau species with various characteristics may co-exist in an AD brain. Here, we directly compared fibrillar SARK and oligomeric HMW tau derived from the same AD brain to determine their relative contribution to tau pathology progression while accounting for interindividual heterogeneity [17]. We show that, despite having similar seeding activities *in vitro* and *in vivo*, only oligomeric HMW tau induces a peculiar rod-like microglial phenotype at the injection site, induces a stronger astrocytic response, has specific effects on tau subcellular localization and possible maturation, and propagates at least as strongly to the entorhinal cortex when injected into the hippocampus of male PS19 mice. Our data strengthen the idea of multiple bioactive tau species within each AD brain and position oligomeric HMW tau as having at least equal potency as SARK tau in terms of seeding and propagation to distally connected regions. If anything, HMW tau appears to generate a more pronounced neuroinflammatory response than fibrillar SARK tau. Since biological potency of the preparations is also likely related to local concentrations, we further postulate that, in the intact brain, the freely soluble HMW species may also be more bioavailable than aggregated tau.

Recent SEC data showed that different tau species can co-exist in the brain ranging from high to low molecular weight species reflecting tau oligomers and tau monomers respectively [57], in addition to the well described fibrillar forms that are the major constituent of NFTs. The fact that HMW tau oligomers are found in low quantities in non-AD brains [57] could reflect either a physiological function for this tau species or reflect a transient state. In contrast, the presence of SARK tau fibrils seems restricted to NFT-bearing cases, suggesting that the presence of fibrillar tau in the brain is abnormal, as no SARK tau fibrils can be isolated from controls. Both SARK and HMW tau have been individually reported in abundance in the AD brain with detectable bioactivities [17, 22, 56, 57]. HMW oligomeric tau is extracted from PBS-soluble brain fractions, while SARK fibrillar tau is extracted from sarkosyl-insoluble brain fractions, suggesting that SARK and HMW tau are two biochemically distinct tau species. Recently, tau filaments have been reported not only in sarkosyl-insoluble AD brain samples but also in aqueous samples [55] and it was shown that short tau filaments are the most seed-competent [28]. However, in our qualitative and quantitative negative stain EM data the low abundance of PHF-like structures in the HMW sample make fibrillar tau unlikely to be responsible for the observed bioactivity of HMW tau. Furthermore, when HMW tau is sonicated 60 pulses similarly to SARK tau, PHF-like structures are not observed anymore but the *in vitro* seeding activity remains further strengthening that the main driving force of the bioactivity of HMW tau is oligomeric tau. In addition, the few PHF-like structures in the HMW sample are similar in size to the PHFs observed in the SARK sample suggesting that fibril size does not account for bioactivity differences. By isolating SARK fibrillar and HMW oligomeric tau from the same AD case, we confirm previous studies [36, 37] showing that two biochemically distinct tau species co-exist in the AD brain.

The development of AD models has allowed the individual study of tau fibrils and tau oligomers which resulted in the conclusion that not only fibrillar [22, 25, 26, 44] but oligomeric tau species too can each be bioactive [7, 13, 35, 43, 45, 53, 62, 66], but no detailed, direct comparison of these two species across multiple cases has been carried out. Using the well-established FRET-based biosensor cell assay [24], we confirm that both SARK tau fibrils and HMW tau oligomers are essentially equally bioactive *in vitro*, suggesting that both have equal ability to recruit the reporter construct, a P301S-mutated tau repeat domain. We next turned to a more complex biological system and took advantage of the commonly used P301S tau transgenic (PS19) mouse model to evaluate the biological activity of the different tau seeds *in vivo*. As expected [8, 17], AD brain derived SARK and HMW tau both increase tau pathology in the hippocampus of PS19 mice 3 months after injection. Consistent with the *in vitro* data, endogenous tau is similarly seeded by SARK fibrillar and HMW oligomeric tau as the number of tau-positive hippocampal neurons is comparable between the two injection groups. These data show again that SARK tau fibrils and HMW tau oligomer preparations, derived from a same brain, have similar seeding abilities, despite exhibiting different biochemical characteristics.

Interestingly, despite similar tau seeding activities between the tau seeds, we observe different tau accumulation patterns after injection into the hippocampus. HMW tau triggers tau pathology in neuronal somas while SARK tau enhances tau accumulation in the dendritic processes of the *stratum oriens* and *stratum radiatum*. We obtain the same results after injecting SARK and HMW tau into the hippocampus of Tau22, mice which express a double-mutated G272V/P301S human tau, suggesting that the neuronal localization of pathological tau protein depends on the characteristics of the injected tau seeds, rather than the nature of the templated endogenous tau protein. These findings echo a recent publication reporting the differential distribution of seeded tau pathology after the injection of PS19 mouse-derived fibrillar and oligomeric tau into 3-month-old PS19 mice [29]. Endogenous tau templated by either tau fibrils or oligomers thus seems to replicate the initial difference between fibrillar and oligomeric tau seeds. However, the mechanisms underlying tau recruitment to the dendritic compartments remain unclear [27].

The hypothesis of prion-like cell-to-cell propagation of tau pathology has been reinforced by several studies in animal models [15, 16]. The injection of AD-derived tau fibrils into the hippocampus of either tau transgenic or WT animals leads to the appearance of tau pathology in brain regions distal but anatomically connected to the injection site [8, 22, 42]. It has also been shown that AD-derived tau oligomers can transmit from cell to cell *in vitro* [57]. We compared the extent of tau pathology in two brain regions distal from the injection site. In this model, while the injection of both SARK tau fibrils and HMW tau oligomers led to similar levels of tau pathology at the injection site in the hippocampus and in the isocortex, and both propagated to peri-/entorhinal cortex, ThioS-positive mature tau aggregates were increased in the synaptically connected peri-/entorhinal cortex in the HMW tau-injected animals only. Similar observations have been made in WT mice, where tau pathology was confined to the injection site after the intrahippocampal injection of brain-derived tau fibrils, but the injection of tau oligomers from the same AD case led to the appearance of tau pathology in distally connected brain regions [36]. This differential ability to propagate between SARK and HMW tau might result from the size of tau particles in each sample. Indeed, studies of the prion protein have shown that the most infectious and propagation-prone particles were about 300-600 kDa, similar size range to HMW tau, compared to larger fibrils [52]. Post-translational modification differences between SARK and HMW tau as reported recently [60] may also account for differential trafficking between these two tau species. Considering the several different pathways of pathological tau secretion and spreading [10] as well as the involvement of neuronal activity in tau pathology progression [12, 46, 63], the rate of progression of SARK and HMW tau could be influenced by multiple, yet unexplored, biological factors.

Besides amyloid plaques and NFTs, AD pathology is accompanied by neuronal and synaptic loss as well as neuroinflammation, including astrocytic and microglial activation [50]. We evaluated changes in glial abundance and morphology in the brain of the injected animals using the commonly used markers GFAP for astrocytes and Iba1 for microglia. Compared to the PBS-injected PS19 mice, only the HMW oligomeric tau-injected animals displayed increased area covered by GFAP suggesting a differential astrocytic response depending on the nature of the injected tau seed. Recent analysis of the astrocytic signature in a P301S tau mouse model showed that reactive astrocytes can adopt multi-faceted changes to adapt to the local environment as shown by increased expression of genes involved in protein degradation for instance [30]. Astrocytic functions involve synaptic pruning [14] and neuronal circuit remodeling [48] as well as neuronal tau uptake [39, 41] making them prone to neurodegeneration and tau propagation. Curiously, we observe increased GFAP staining in the HMW tau-injected animals, which are the only ones showing tau propagation to the entorhinal cortex.

On the other hand, we did not find any group difference in Iba1 percent area at the injection site, despite them being different in the amount of tau pathology, suggesting that Iba1-related microglial response is not affected by the increase of tau-positive neurons in this model. The PS19 mouse model exhibits signs of neuroinflammation beginning at 3 months of age, before the appearance of tau aggregates [66]. Therefore, the absence of a difference between the tau-injected and control animals at the time of sacrifice (6 months of age) suggests that microglial activation might have plateaued when the first spontaneous or induced tau tangles appeared. These data are consistent with a previous study, where 6 months after the injection of AD brain homogenates into ALZ17 Tau transgenic mice, no difference in microglial activation was observed using the Iba1 microglial marker compared to controls [15].

Surprisingly, when we looked closer at the morphology of Iba1-positive microglia, we observed the appearance of abundant rod-like microglia in the hippocampus of HMW tau-injected animals only. Rod microglia were first described a century ago by Nissl, yet their function and origin remain unknown. To this day, rod microglia are only characterized by their morphology presenting an elongated and narrow cell body and very few planal processes. These rod microglia are often described as following and enveloping neuronal processes, and can also assemble forming long train-like structures [21]. Historically, rod microglia were described in the brain of patients suffering from an infectious disease like typhus or syphilis [58]. More recently, rod microglia have been reported in the brain of elderly [5], AD patients [6], in cases of traumatic brain injury (TBI) [61, 68], epilepsy [64] and in hypoxic/ischemic conditions [34]. Interestingly, these Iba1-positive rod microglia that we solely observe in the HMW tau-injected PS19 mice are also present in the grey matter of the AD cases from where the tau seeds were extracted, showing that the injection model recapitulates the human AD pathological environment and suggesting that in the human AD brain rod microglia might be somehow triggered by oligomeric forms of tau.

Very little is known about this microglial phenotype, therefore we explored the expression of Clec7a, one marker of disease-associated (DAM) microglia together with other markers [31] at the crosstalk between infectious diseases and AD. Clec7a (or Dectin-1) is a β-glucan receptor expressed by myeloid cells with both exogenous microbial and endogenous ligands [40]. Interestingly, Clec7a has been reported to be involved in the inflammatory response following not only AD but also TBI and stroke [65], all being pathological conditions associated with rod microglia. Triple staining for microglia (Iba1), pathological tau (AT100) and Clec7a in the different conditions revealed that Clec7a-positive Iba1-positive microglia are restricted to the HMW oligomeric tau-injected animals, and that these cells are not necessarily in the vicinity of tau-positive somas or processes. In the hippocampus, all Clec7a-positive microglia seem rod-like, but not all rod-like microglia are Clec7a-positive, suggesting an additional molecular level of diversity among morphologically similar microglia. In this model, the presence of rod microglia is restricted to the injection site, suggesting the involvement of the injected HMW oligomeric tau in the trigger of this peculiar microglial phenotype. In contrast, we observed Clec7a-positive microglia all along the propagation pathway of bioactive tau seeds, starting from the targeted CA1 pyramidal neurons to the entorhinal cortex. Clec7a-positive microglia were particularly abundant in the *stratum radiatum* and the molecular layer of the hippocampus where CA1 pyramidal dendrites and axons are located, respectively, suggesting that Clec7a-positive microglia could be a readout for tau seed propagation.

In summary, we show that SARK fibrillar and HMW oligomeric tau samples derived from the same AD case induce similar tau related phenotypes yet trigger differential and specific cellular effects. Both tau fibrils and tau oligomers can seed endogenous tau, but patterns of neuronal tau accumulation differ, as do the properties of propagated species. Most importantly, we report a differential glial response with the appearance of a peculiar and poorly characterized rod-like microglial phenotype in the oligomeric tau-injected animals only. Therefore, the data strongly suggest that various tau seeds co-exist in the AD brain and that these tau seeds have unique intrinsic properties that are, in the present study, reflected at the biological level. These observations confirm the intraindividual heterogeneity of tau species in AD, opening doors to novel and better targeted anti-tau therapeutic strategies.

## Acknowledgements

We are grateful for the AD patients and the Massachusetts ADRC brain bank (P30AG062421) for providing the tissue samples. The current work was funded by NIH RF1AG058674, RF1AG059789, R56AG061196, and support from the Rainwater Foundation, Cure Alzheimer Fund, and the JPB foundation.

## Authors contributions

AMG and BTH conceived the study and designed the experiments. AMG, LAW, AK, CC, CG and JEC performed experiments and analyzed data. RP provided critical input on the study. The manuscript was written by AMG and BTH with input from all the authors.

