## Supplementary figures for "Tau seeding and spreading in vivo is supported by both AD-derived fibrillar and oligomeric tau"

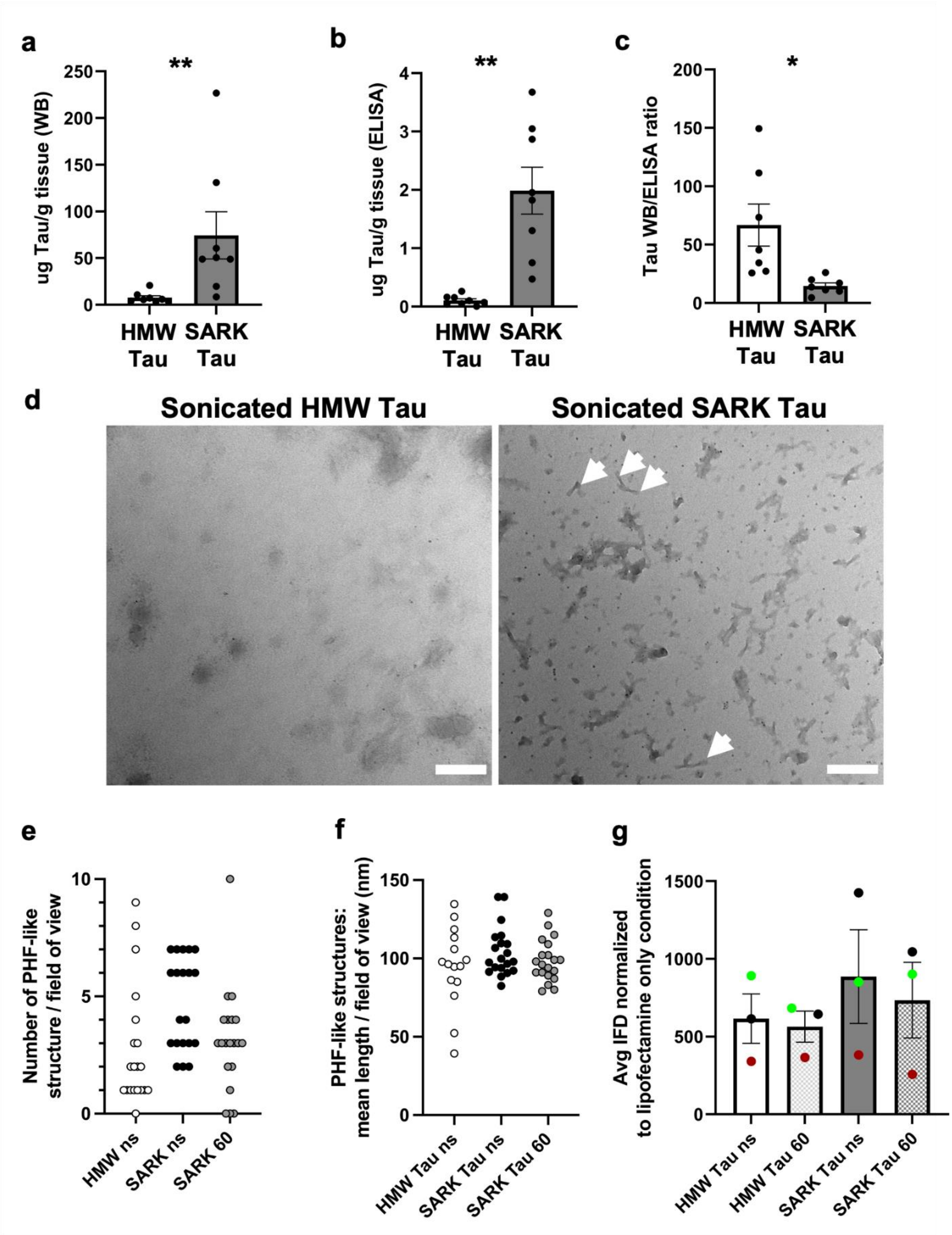

**Fig. S1: Tau fibrils are more abundant than HMW tau oligomers per tissue weight**

**a.** Quantification of total tau extracted with each method by denaturing WB using the polyclonal tau DAKO antibody. **b.** Quantification of total tau extracted with each method by non-denaturing tau ELISA. **c.** WB/ELISA ratio reflecting tau oligomerization in each tau extract. Data represented as mean±SEM, Wilcoxon matched-pairs test, *$p<0.05$, **$p<0.01$. **d.** Representative negative stain EM images of sonicated (60 short pulses) samples from AD case #2399 used for *in vivo* injections. Sonicated HMW tau does not display any PHFs while PHFs are still abundant in the SARK tau sample after sonication. Scale bar = 200 nm. **e-f.** Quantification of the number of PHF-like structures and the fibril length in 20 randomly sampled negative stain EM fields of view for each tau sample from AD case #2399 used for *in vivo* injections. Similar results were obtained for 4 separate AD cases, suggesting that these data are representative of AD preparations. **g.** Quantification of the seeding activity in non-sonicated and sonicated HMW and SARK samples by flow cytometry 24 hours after transfection. Integrated FRET densities were normalized to the lipofectamine only negative control. Data represented as mean±SEM from 3 independent experiments, one color per AD case, Friedman test for paired data, non-significant. ns=non-sonicated; 60=sonicated 60 pulses.

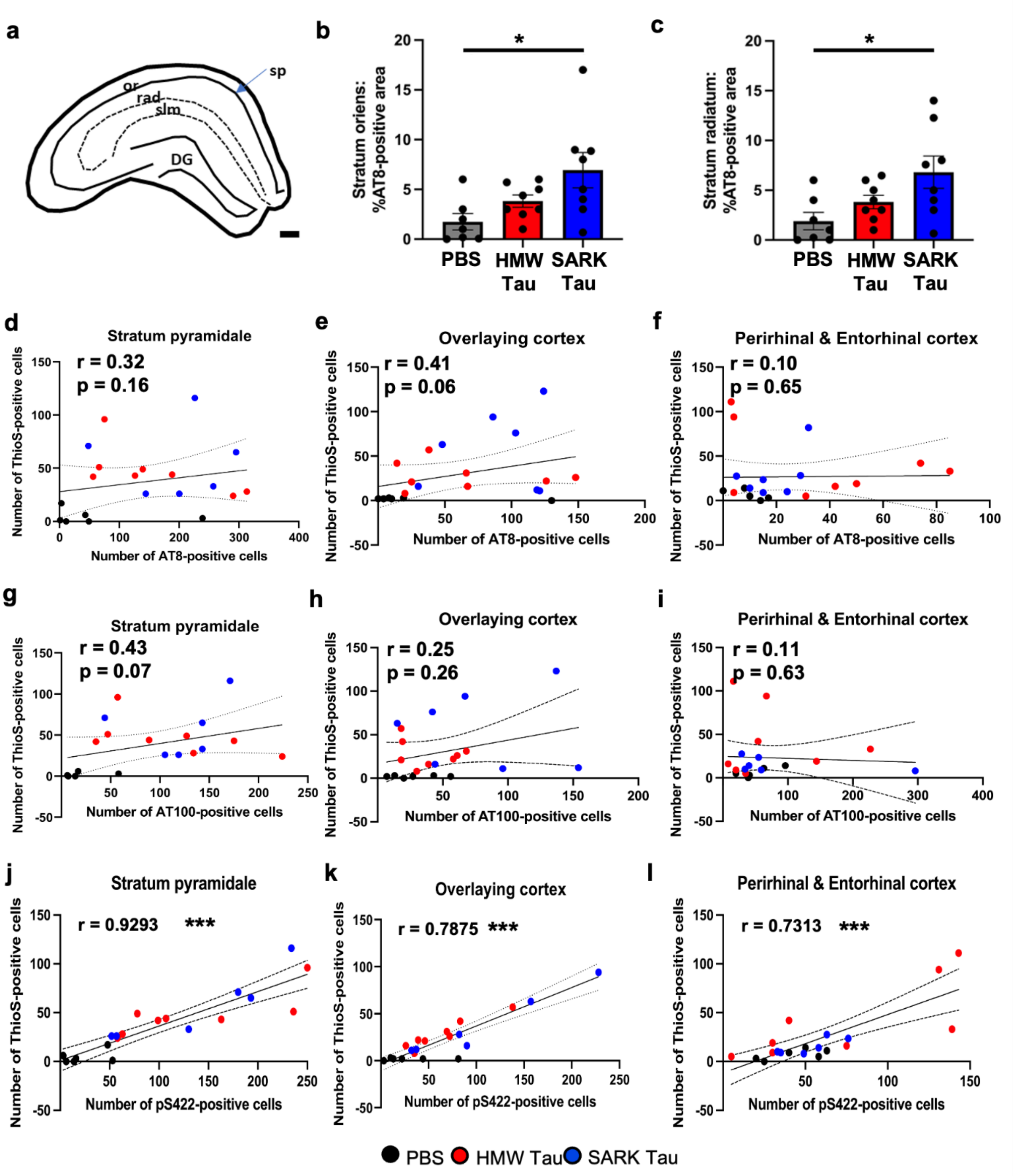


**Fig. S2: Tau aggregation does not correlate with AT8 or AT100 phosphorylation**

**a.** Schematic of the dorsal hippocampal layers. or: *stratum oriens*, sp: *stratum pyramidale*, rad: *stratum radiatum*, slm: *stratum lacunosum moleculare*, DG: dentate gyrus. Scale bar = 20 µm. **b.** Quantification of the percentage of the AT8-positive area in the *stratum oriens* of the dorsal hippocampus of PS19 mice 3 months after injection. **c.** Quantification of the percentage of the

AT8-positive area in the *stratum radiatum* of the dorsal hippocampus of PS19 mice 3 months after injection. Data represented as mean±SEM, Kruskal-Wallis, Dunn's multiple comparison, *<p0.05. **d-f.** Correlation analysis between the number of ThioS-positive cells and the number of AT8-positive cells in the dorsal hippocampus, the overlying, and the peri-/ento-rhinal cortices respectively in PS19 mice 3 months after injection. Data represented with the 95% confidence interval, Spearman correlation, not significant. **g-i.** Correlation analysis between the number of ThioS-positive cells and the number of AT100-positive cells in the dorsal hippocampus, the overlying, and the peri-/ento-rhinal cortices respectively in PS19 mice 3 months after injection. **j-l.** Correlation analysis between the number of ThioS-positive cells and the number of late stage phosphorylated pS422-positive cells in the dorsal hippocampus, the isocortex and the peri-/ento-rhinal cortices respectively in PS19 mice 3 months after injection. Data represented with the 95% confidence interval, Spearman correlation, ***p<0.001.

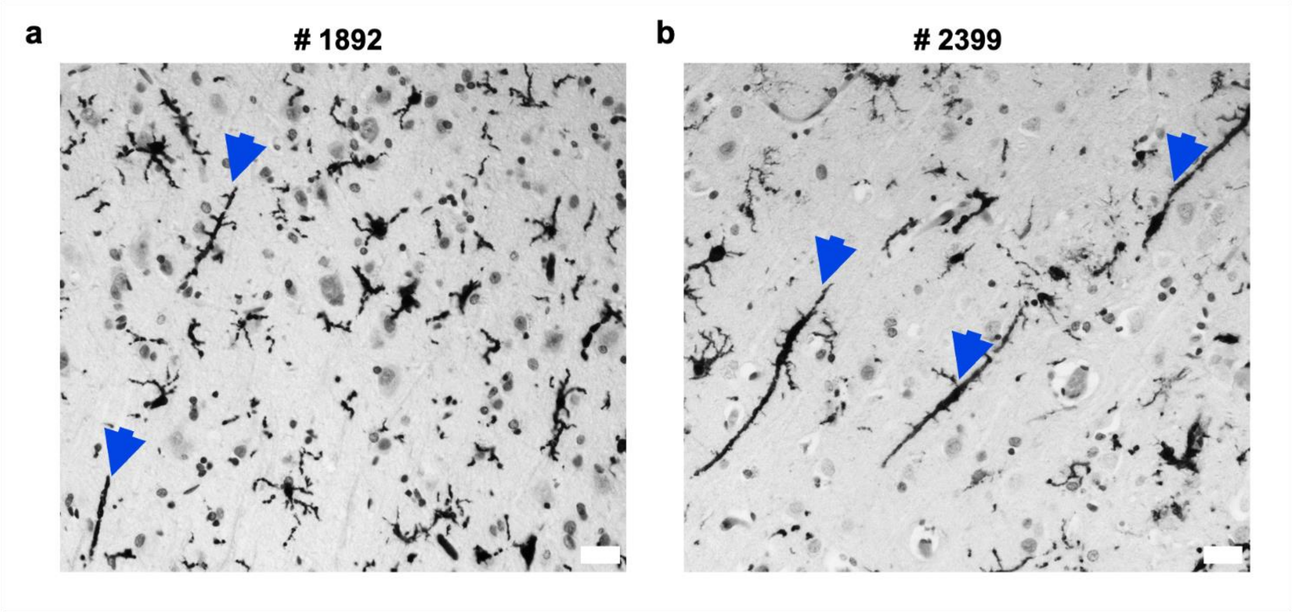


**Fig. S3: Rod microglia is present in the AD cases used for the *in vivo* studies**

**a-b.** Representative images of Iba1-positive rod microglia (blue arrows) in the grey matter of the parietal cortex (BA 7) of AD case #1892 and #2399, respectively. The injection of HMW tau derived from the parietal grey matter of these two cases into PS19 mice led to the appearance of rod microglia at the injection site thus recapitulating the AD pathological environment. Scale bars = 20 µm.

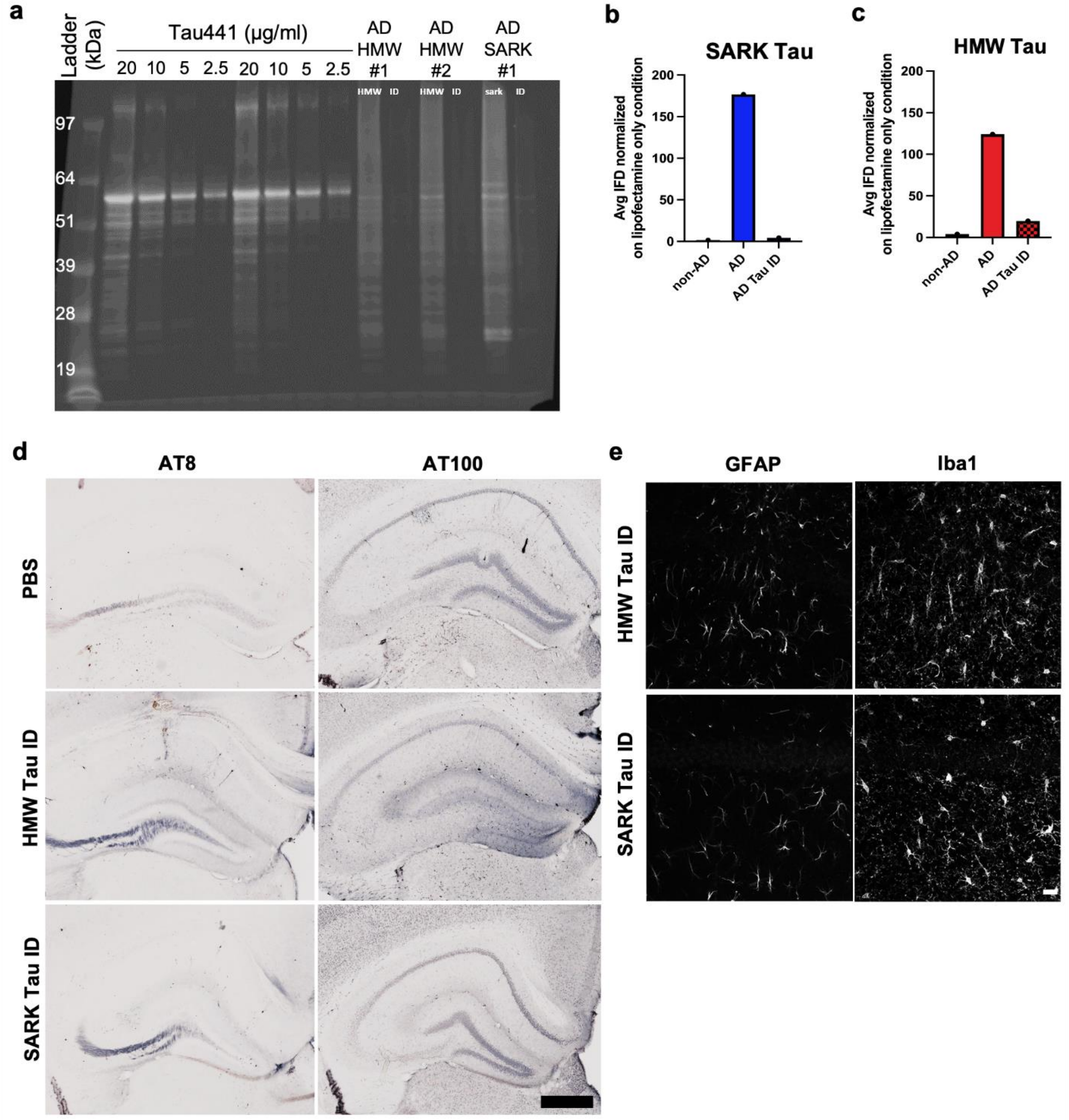


**Fig. S4: Confirmation of the tau specificity of the effect**

**a.** Total tau WB (DAKO) after immunodepletion of tau with HT7 antibody in both SARK and HMW tau samples. **b-c.** *In vitro* seeding activity normalized to the lipofectamine only negative control of SARK and HMW tau respectively. Immunodepletion of tau with HT7 antibody greatly reduces the seeding activity of both SARK and HMW tau samples to a level similar to a non-AD control. **d.** Representative images of AT8- and AT100- tau pathology in HT7 tau-immunodepleted

SARK and HMW tau-injected PS19 mice 3 months after injection. Scale bars = 500 µm. **e.** Representative images of GFAP and Iba1 glial stainings in HT7 tau-immunodepleted SARK and HMW tau-injected PS19 mice 3 months after injection. Scale bars = 20 µm. ID = immunodepleted.

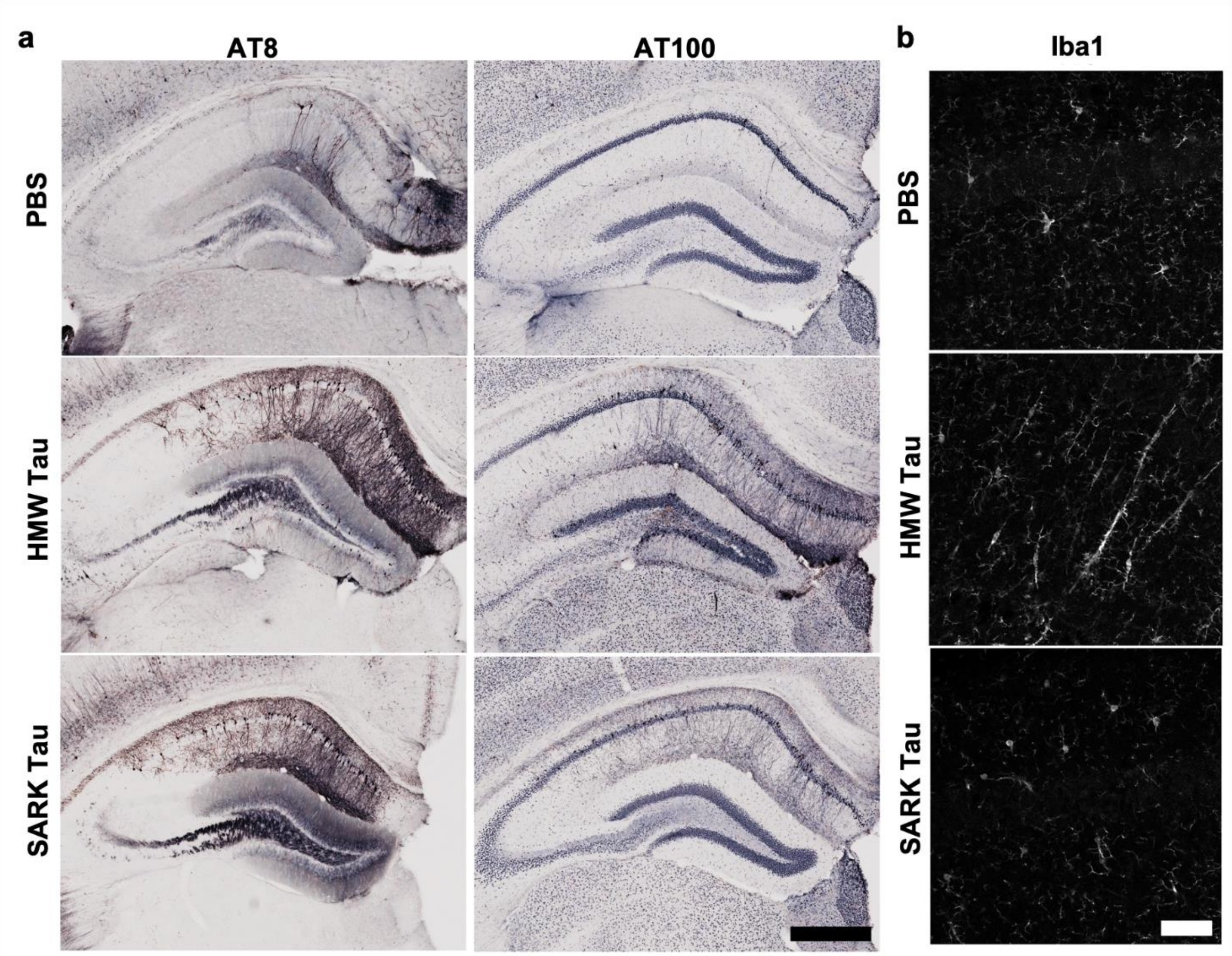


**Fig. S5: Tau oligomers and tau fibrils behave similarly whatever tau is present in the recipient cell**

**a.** Representative images of phosphorylated AT8- and AT100- tau pathology in the dorsal hippocampus of Tau22 mice 2 months after injection. Scale bar = 500 µm. **b.** Representative images of Iba1-positive microglia in the dorsal hippocampus of Tau22 mice 2 months after injection. Rod-like microglia is only observed in the HMW tau-injected group. Scale bar = 50 µm.
