## Supplementary table for "Tau seeding and spreading in vivo is supported by both AD-derived fibrillar and oligomeric tau"

| Antibody | Reactivity | Species | Reference | IHC dilution | IF dilution | WB dilution |
|---|---|---|---|---|---|---|
| **AT8 (biotinylated)** | Human Tau pS212, pT214 | Mouse | Thermofisher (MN1020B) | 1:400 | 1:400 | |
| **AT100** | Human Tau pS202, pT205 | Mouse | Thermofisher (MN1060) | 1:4000 | 1:1000 | |
| **Bassoon** | Mouse presynaptic | Guinea pig | Synaptic systems (141 004) | | 1:1500 | |
| **Clec7a (murine)** | murine Dectin-1 | Rat | InvivoGen | | 1:50 | |
| **GFAP-CY3** | GFAP | Mouse | Sigma-Aldrich (C9205) | | 1:10,000 | |
| **Iba1** | Iba1 | Rabbit | Wako (019-19741) | | 1:1000 | |
| **Iba1** | Iba1 | Rabbit | Abcam (ab178847) | 1:500 (autostainer) | | |
| **pS422** | Human Tau pS422 | Rabbit | Abcam (ab79415) | | 1:1000 | |
| **PSD95** | Mouse postsynaptic | Rabbit | Thermofisher (51-6900) | | 1:1000 | |
| **Tau DAKO** | Total Tau | Rabbit | DAKO (A0024) | | | 1:5000 |

**Table** S1: list of the primary antibodies used in the present study and their working concentration.
